# A perturbed gene network containing PI3K/AKT, RAS/ERK, WNT/β-catenin pathways in leukocytes is linked to ASD genetics and symptom severity

**DOI:** 10.1101/435917

**Authors:** Vahid H. Gazestani, Tiziano Pramparo, Srinivasa Nalabolu, Benjamin P. Kellman, Sarah Murray, Linda Lopez, Karen Pierce, Eric Courchesne, Nathan E. Lewis

**Affiliations:** Autism Center of Excellence, Department of Neuroscience, University of California San Diego, La Jolla, California, USA; Department of Pediatrics, University of California San Diego, La Jolla, California, USA; Novo Nordisk Foundation Center for Biosustainability, University of California San Diego, La Jolla, California, USA; Bioinformatics and Systems Biology Program, University of California San Diego, La Jolla, California, USA; Department of Pathology, University of California San Diego, La Jolla, California, USA; Department of Bioengineering, University of California San Diego, La Jolla, California, USA

## Abstract

Hundreds of genes are implicated in autism spectrum disorder (ASD) but the mechanisms through which they contribute to ASD pathophysiology remain elusive. Here, we analyzed leukocyte transcriptomics from 1-4 year-old male toddlers with ASD or typical development from the general population. We discovered a perturbed gene network that includes genes that are highly expressed during fetal brain development and which is dysregulated in hiPSC-derived neuron models of ASD. High-confidence ASD risk genes emerge as upstream regulators of the network, and many risk genes may impact the network by modulating RAS/ERK, PI3K/AKT, and WNT/β-catenin signaling pathways. We found that the degree of dysregulation in this network correlated with the severity of ASD symptoms in the toddlers. These results demonstrate how the heterogeneous genetics of ASD may dysregulate a core network to influence brain development at prenatal and very early postnatal ages and, thereby, the severity of later ASD symptoms.

## INTRODUCTION

Autism spectrum disorder (ASD) is a neurodevelopmental disorder with prenatal and early postnatal biological onset^1–3^. Genetic factors contribute to the predisposition and development of ASD with estimated heritability rates of 50-83%^4, 5^. Large genetic studies have implicated several hundred ASD risk (rASD) genes that could be associated with many different pathways, cellular processes, and neurodevelopmental stages^6–10^. This highly heterogeneous genetic landscape has raised challenges in elucidating the biological mechanisms involved in the disorder. While rigorous proof remains lacking, current evidence suggests that rASD genes fall into networks and biological processes^6, 7, 9, 11^ that modulate one or more stages of prenatal and early postnatal brain development, including neuron proliferation, migration, neurite growth, and synapse formation or function^3, 8^. However, these insights are mostly gained from studies focused on single rASD genes^3^ or transcriptome analyses of neurotypical brains^9, 11^. Thus, we have an incomplete picture of the molecular changes at the individual level and their role in the observed heterogeneity in the core clinical deficits of socialization and restricted, repetitive behavior.

To further complicate efforts, rASD genes have been largely identified through *de novo* loss-of-function mutations in their coding sequence. Such events contribute to ASD risk in 5-10% of the ASD population, and most of heritable genetic risk is thought to reside in common variants that are also found in subjects with typical development^5, 12–15^. It remains unclear whether ASD in subjects with known rASD gene mutations manifests as ASD subtypes with distinct molecular etiologies, or whether the underlying mechanisms are shared with the overall population of subjects with ASD.

To address these questions, it is important to understand which molecular processes are perturbed in prenatal and early postnatal life in individuals with ASD, assess how they vary among subjects, and evaluate how these perturbations relate to rASD genes and early-age clinical ASD symptoms. It is expected that the genetic changes in ASD alter gene expression and signaling in the developing brain^3, 7, 10^. Therefore, capturing dysregulated gene expression at prenatal and early postnatal ages may unravel the molecular organization underlying ASD. Unfortunately, doing so is challenging as ASD cannot be clinically diagnosed at these early stages based on currently established behavioral symptoms^16^, and accessible postmortem brain tissues are from much older individuals with ASD, well beyond the ages when rASD genes are at peak expression and long after ASD diagnosis. In contrast to living neurons, which have a limited time window for proliferation and maturation, other cell types constantly regenerate, such as blood cells. Given the strong genetic basis of ASD, some dysregulated developmental signals may continually reoccur in blood cells and thus be studied postnatally^17–20^.

Reinforcing this notion, it was recently demonstrated that genes that are broadly expressed across many tissues are major contributors to the overall heritability of complex traits, and it was postulated that this could be relevant to ASD^21^. Lending credence to this, previous studies have reported that differentially expressed genes in the blood of subjects with ASD are enriched for regulatory targets of *CHD*8^18^ and *FMR1*^22^, two well-known rASD genes. Similarly, both lymphoblastoid cells of subjects with ASD and hiPSC-derived models of fragile-X syndrome show over-expression of mir-181 with a potential role in the disorder^23^. Likewise, leukocytes from toddlers with ASD show perturbations in biological processes such as cell proliferation, differentiation, and microtubules^24–27^, and these coincide with dysregulated processes seen in hiPSC-derived neural progenitors and neurons from individuals with ASD and brain enlargement^28, 29^. Ultimately, establishing the signatures of ASD in other tissues will facilitate the study of the molecular basis of the disorder in the first few years of life.

Here we leverage transcriptomic data from leukocytes, hiPSC-derived neuron models, and the neurotypical brain to study the architecture of transcriptional dysregulation in ASD, its connection to rASD genes, and its association with prenatal brain development and postnatal socialization symptom severity in ASD. We discovered a conserved dysregulated gene network by analyzing leukocyte transcriptomic data from 1-4 year-old toddlers with ASD and typical development (TD). The dysregulated network is enriched for pathways known to be perturbed in ASD neurons, involves genes that are highly expressed in prenatal brain, and is dysregulated in hiPSC-derived neurons from subjects with ASD and brain enlargement. Consistent with the postulated structure of complex traits^21, 30^, we show that rASD genes in diverse functional groups converge upon and regulate this core network. Importantly, we found the dysregulation extent of this core network is correlated with the severity of socialization deficits in toddlers with ASD. Thus, our results demonstrate how the heterogeneous rASD genes converge and regulate a biologically relevant core network, capturing the possible molecular basis of ASD.

## RESULTS

### Increased transcriptional activity in leukocytes from toddlers with ASD

We analyzed leukocyte gene expression profiles obtained from 226 male toddlers (119 ASD and 107 TD). Robust linear regression modeling of the data identified 1236 differentially expressed (DE) genes (437 downregulated and 799 upregulated; FDR <0.05). Jack-knife resampling demonstrated that the expression pattern of DE genes was not driven by a small number of cases, but rather shared between the vast majority of subjects with ASD (Fig. S1). We further validated the expression patterns in additional replicate and independent cohorts (Fig. S1-S4).

In many disease conditions, transcriptional programs in cells deviate from normal states due to dysregulations in signaling pathways, transcription factors and epigenetic marks. Therefore, we employed a systems approach to decipher network-level transcriptional perturbations in leukocytes of toddlers with ASD (Fig. 1). We reasoned that perturbations to ASD-associated molecular pathways would be reflected in the co-expression patterns between DE genes. To identify such ASD-relevant dysregulations, we first extracted a static gene network (that is, the network is indifferent to the cell context) composed of all known high-confidence physical and regulatory interactions among the DE genes (Methods). We next pruned the static network using our leukocyte transcriptome data to obtain context-specific networks of each diagnosis group separately (that is, the networks differ in genes and their interactions, based on their associated gene expression data). Specifically, context-specific networks were built for each of ASD and TD groups by only retaining those interactions from the static network that were significantly co-expressed (FDR <0.05) within the group. Both context-specific networks, called DE-ASD and DE-TD, were constructed based on the same static network from the same set of genes (i.e., genes that are expressed in leukocytes and show differential expression). However, following removal of interactions lacking co-expression, a proportion of genes become unconnected and these were consequently removed from the DE-ASD and DE-TD networks. Therefore, DE-ASD and DE-TD exhibited 63% overlap in their gene composition, with differences mostly related to genes that were loosely connected in the starting static network. To ensure the robustness of our conclusions to the changes in the structure of the static network, we replicated all presented results on two other static networks with higher density of interactions that resulted in a higher number of overlapping genes between corresponding context-specific networks (Methods).

**Fig 1.**
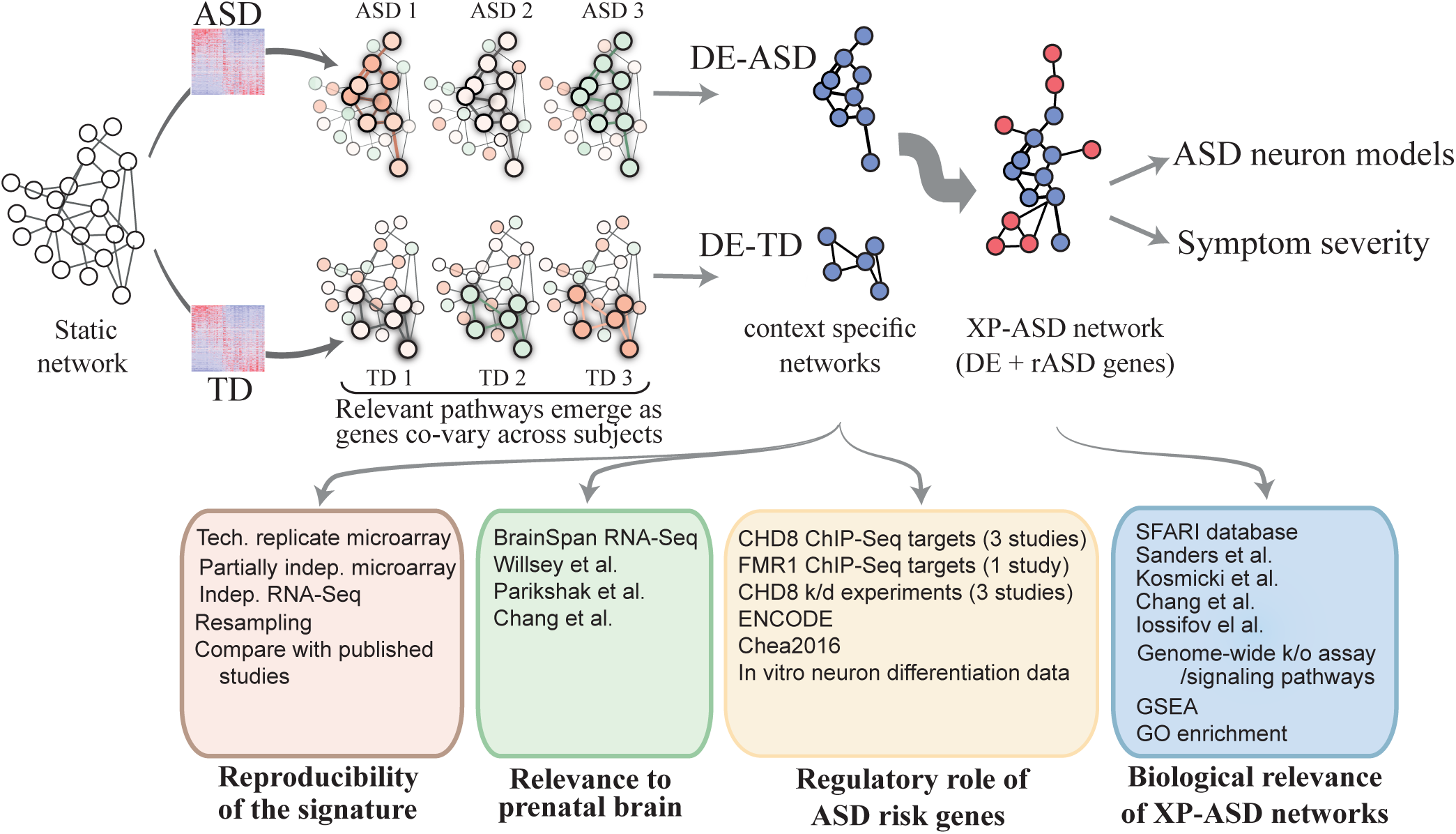
Study overview. Transcriptome analysis of 226 toddlers with ASD or typical development identified 1236 DE genes. We built a comprehensive “static” network of DE genes from high confidence physical and regulatory interactions from the Pathway Commons, BioGrid, and Reactome databases. To identify transcriptional programs that are active in each diagnosis group, we retained pairs of interacting genes in the static network that are highly co-expressed in each diagnosis group. This yielded context specific DE-ASD and DE-TD networks, allowing to compare the activity of transcriptional programs between ASD and TD conditions. To connect the DE-ASD network to ASD risk genes, an XP-ASD network was built using DE and ASD risk (rASD) genes. The DE-ASD and XP-ASD networks were analyzed in the context of neural differentiation, ASD neuron models, and ASD symptom severity. To ensure results were robust to variations in the interaction networks, we reproduced the results by replacing the high confidence static network (the first step in pipeline) with a functional and a full co-expression network (Methods).

To test if transcriptional programs were being modulated in ASD, we merged the genes and interactions in the DE-ASD and DE-TD networks, and compared the ‘co-expression magnitude’ of interactions in the merged network between ASD and TD samples^31–33^. This proxy for the transcriptional activity of gene networks^9^ demonstrated that co-expression magnitude was higher in the ASD than the TD samples (Fig. 2a; p-value <0.01; paired Wilcoxon-Mann-Whitney test). The stronger co-expression in the DE-ASD network suggests a higher level of concerted activation or suppression of pathways involving DE genes among the subjects with ASD. Further analysis confirmed that the changes in the co-expression magnitude, rather than the gene composition, is the primary driver of the elevated network transcriptional activity. This higher level of concerted co-regulation of the network was also reproducible in two additional ASD transcriptomic datasets and across alternative analysis methods (Fig. S1-S4).

**Fig 2.**
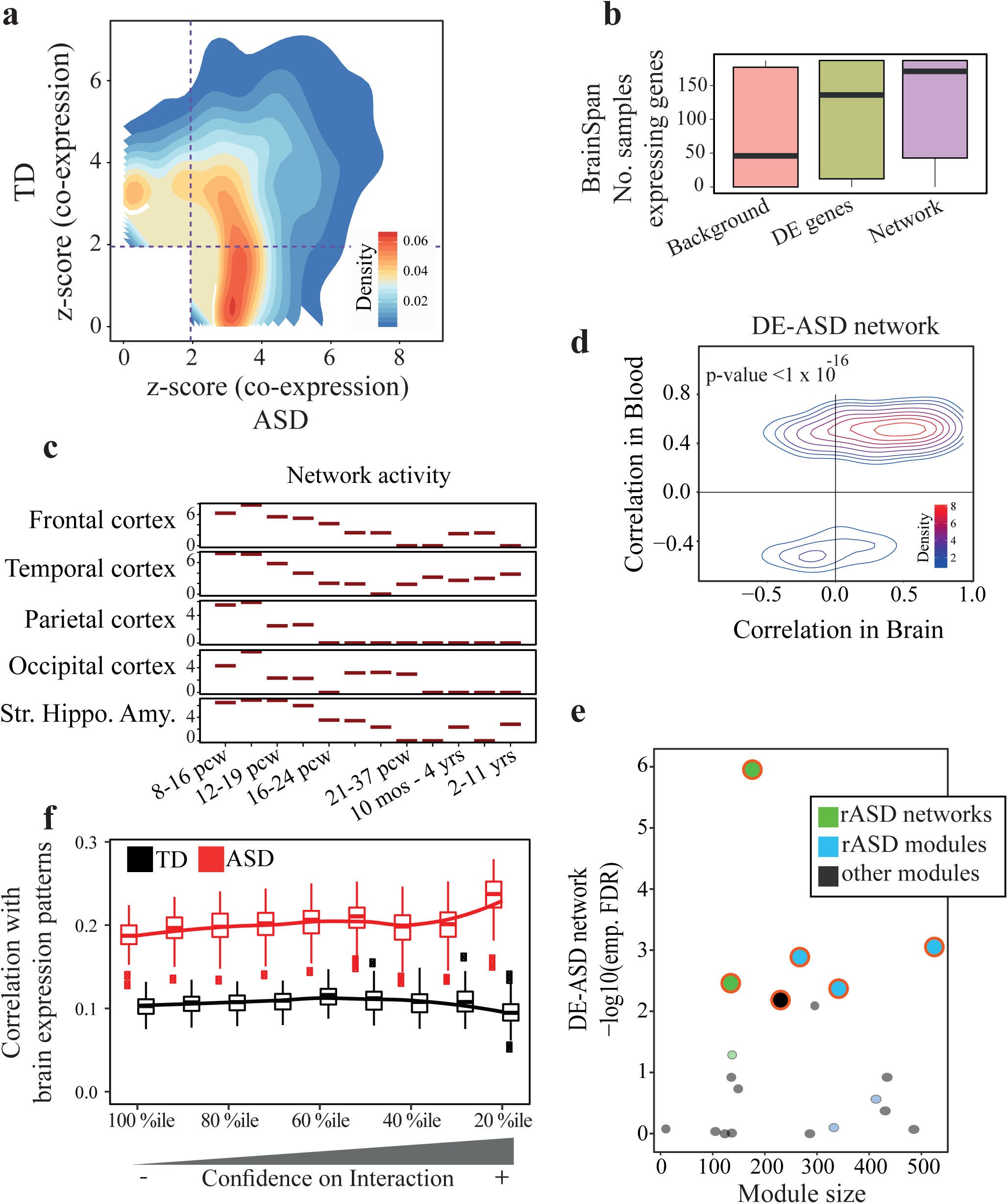
Co-expression activity is elevated in the DE-ASD network in ASD leukocytes and preserved in prenatal brain. a) The DE-ASD network shows stronger co-expression in ASD toddlers compared to TD toddlers, suggesting pathways in the DE-ASD network are being modulated in ASD. For an unbiased analysis, the union of genes and interactions from DE-ASD and DE-TD networks was considered for this analysis (n= 119 ASD and 107 TD toddlers; see also Fig. S3). b) Genes in the DE-ASD network are highly expressed in the brain between 8 post conception weeks (pcw) to 1 year-old. For each gene, samples strongly expressing the gene (RPKM >5) were counted, based on BrainSpan normalized RNA-Seq data^34^. The background genes included all protein coding genes expressed in our microarray experiment and present in BrainSpan (n= 187 neocortex samples; see also Fig. S5). c) The activity pattern of the DE-ASD network across brain regions during neurodevelopment. At each time window, the distribution of co-expression magnitudes of interacting gene pairs in the DE-ASD network was measured using unsigned Pearson’s correlation coefficient (n= 121 frontal, 73 temporal, 42 parietal, 27 occipital cortices, and 72 striatum, hippocampus, and amygdala samples across time points). The co-expression values were next compared to a background distribution using a Wilcoxon-Mann-Whitney test (Methods). The y-axis shows z-transformed p-values of this comparison. d) Leukocyte gene co-expression in the DE-ASD network is conserved in the prenatal and early postnatal neocortex transcriptome. The Pearson’s correlation coefficient of interacting gene pairs in the DE-ASD network was calculated from the neocortex transcriptome (n= 187 neocortex samples; 8 pcw until 1 year-old). The correlations were next paired with those in ASD group (n=119 subjects). A p-value was estimated by comparing the observed preservation of DE-ASD with that of DE-TD using a re-sampling method (Fig. S6). e) Overlaps of the DE-ASD network with brain developmental modules and networks. Modules and networks enriched for rASD genes significantly overlap with the DE-ASD network (FDR <0.1; permutation test). rASD networks: networks constructed around high confidence rASD genes^7, 9^; rASD modules: co-expression modules enriched for rASD genes^11^; other modules: modules that are not enriched for rASD genes^11^. f) Similarity of interactions of a brain co-expression network around rASD genes^9^ with ASD and TD samples as measured by Pearson’s correlation coefficient. Boxplots represent the similarity based on 100 random sub-samplings (n=75 ASD and 75 TD). The x-axis represents the top percentile of positive and negative interactions based on the brain transcriptome interaction correlation value. Brain co-expression is based on transcriptome data from 10–19 pcw (see also Fig S5). Boxplots represent the median (horizontal line), lower and upper quartile values (box), and the range of values (whisker), and outliers (dots).

In summary, the leukocyte transcriptional networks of the DE genes show higher than normal co-expression activity in ASD. Moreover, the dysregulation pattern is present in a large percentage of toddlers with ASD, as evidenced by the resampling analyses and the other two ASD datasets.

### The leukocyte-based gene network captures transcriptional programs associated with brain development

We next assessed the potential involvement of the leukocyte-based network to gene expression patterns during brain development. By overlaying the neurodevelopmental RNA-Seq data from BrainSpan^34, 35^ on our DE-ASD network, we found that the DE-ASD network was enriched for highly expressed genes in the neocortex at prenatal and early postnatal periods (p-value <4.3×10^-30^; Fig. 2b).

To investigate the spatiotemporal activity of the DE-ASD network during brain development, we measured the magnitude of gene co-expression within the DE-ASD network at different neurodevelopmental time windows across brain regions. We found that the highest levels of co-expression of the DE-ASD network temporally coincided with peak neural proliferation in brain development (10-19 post conception weeks^3, 8^), after which co-expression activity gradually decreased (Fig. 2c; Fig. S5). Expression levels of genes in the DE-ASD network followed a similar pattern (Fig. S6). Further supporting the transcriptional activity of the leukocyte-derived DE-ASD network in prenatal brain, we found evidence that the network is mostly preserved at the co-expression level between ASD leukocytes and prenatal brain. Specifically, the direction of correlations (i.e., positive or negative) in the leukocyte transcriptome of subjects with ASD is mostly preserved in prenatal and early postnatal brain (Fig. 2d). Importantly, this preservation of co-expression was significantly higher in the DE-ASD network than in the DE-TD network (p-value <10^-16^; Fig. S6).

### rASD genes are associated with the DE-ASD network

We next analyzed the DE-ASD network in the context of other studies to test the relevance of our DE-ASD network to ASD. Parikshak et al. previously reported gene co-expression modules associated with cortical laminae development during prenatal and early postnatal ages^11^. A subset of these modules show enrichment in rASD genes^11^. We examined the overlap of our leukocyte-derived network with all modules from Parikshak et al^11^. The DE-ASD network preferentially overlapped with rASD gene-enriched modules from that study (Fig. 2e). This suggests that our DE-ASD network is functionally related to rASD genes during neocortical development. Our DE-ASD network also overlapped with the networks of rASD genes reported in other studies^7, 9^, indicating the robustness of the results (Fig. 2e). Intriguingly, the prenatal brain co-expression network of high-confidence rASD genes was more similar to that of ASD leukocytes than TD leukocytes (Fig. 2f), suggesting that neurodevelopmental transcriptional programs related to rASD genes might be more active in the leukocyte transcriptome of toddlers with ASD than in that of TD toddlers.

With the observed overlap patterns, we next tested for enrichment of rASD genes in the DE-ASD network. For this analysis, we considered different rASD gene lists of different size and varying confidence levels (Methods). Surprisingly, this analysis demonstrated that rASD genes are not enriched in the DE-ASD network (p-value >0.19).

### The DE-ASD network is enriched for regulatory targets of rASD genes

Many high confidence rASD genes have regulatory functions^3, 7, 10^. Although the perturbed DE-ASD network is not enriched for rASD genes, it overlaps with brain co-expression modules and networks containing known rASD genes. At the mechanistic level, the observed co-expression of rASD and DE genes in the prenatal brain could be due to the regulatory influence of rASD genes on the DE-ASD network, and thereby genetic alterations in rASD genes could cause the transcriptional perturbation and the increase in gene co-expression within the DE-ASD network.

To elucidate if rASD genes could regulate the DE-ASD network, we examined if the regulatory targets of rASD genes are enriched in the DE-ASD network. Indeed, we observed that the DE-ASD network is enriched for genes regulated by two high-confidence rASD genes, *CHD8*^36–38^ and *FMR1*^39^ (Fig. 3a). To more systematically identify regulators of the network, we evaluated the overlap of the DE-ASD network with the regulatory targets of rASD transcription factors from the ENCODE project^40^ and Chea2016 resource^41^. Strikingly, the DE-ASD network is significantly enriched for the regulatory targets of 11 out of 20 high-confidence, strong-candidate and suggestive-evidence rASD genes (SFARI categories 1-3) (OR: 2.54; p-value: 0.05; Fig. 3b).

**Fig 3.**
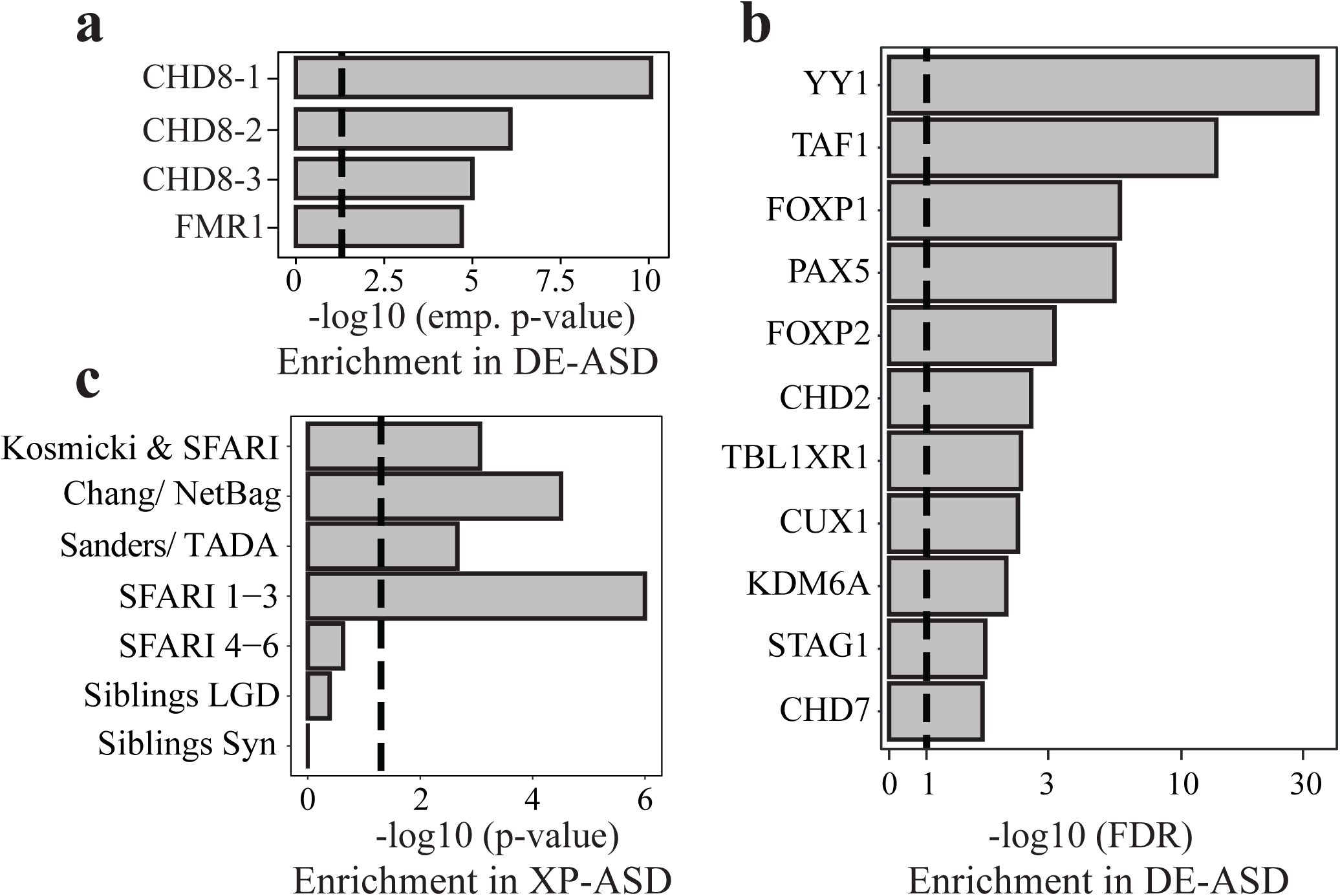
rASD genes are enriched for the regulators of the DE-ASD network. a) Genes identified by ChIP-Seq as regulatory targets of *CHD8* (*CHD8*-1: Sugathan et al.^36^; *CHD8*-2: Gompers et al.^38^; *CHD8*-3: Cotney et al.^37^) and *FMR1*^39^, two high confidence rASD genes, are enriched in the DE-ASD network. Enrichment was assessed empirically (see also Fig. S7); dashed line shows p-value 0.05. b) The DE-ASD network significantly overlaps with the regulatory targets of rASD genes based on the ENCODE and Chea2016 repositories (FDR <0.1; hypergeometric test); dashed line shows FDR 0.1. c) High confidence genes are significantly enriched in the XP-ASD network (hypergeometric test). The lists of high confidence rASD genes were extracted from SFARI database^42^, Kosmicki et al.^14^, Chang et al.^7^, and Sanders et al.^15^. List of likely gene damaging (LGD) and synonymous (Syn) mutations in siblings of ASD subjects were extracted from Iossifov et al.^13^ Dashed line indicates p-value 0.05.

### The DE-ASD network is preferentially linked to high-confidence rASD genes

rASD genes were often not differentially expressed in ASD leukocytes, and the DE-ASD network was therefore not enriched in rASD genes. To explore if rASD genes may nevertheless regulate the DE-ASD network, we expanded the DE-ASD network by including rASD genes. Thus, we obtained an expanded-ASD (XP-ASD) network. To construct the XP-ASD network, we used a similar approach to that used for the DE-ASD network. Briefly, we built a high-confidence static network of DE and 965 candidate rASD genes. The context-specific XP-ASD network was next inferred by retaining only the significantly co-expressed interacting pairs in ASD samples. This pruning step removed genes from the static network that were not significantly co-expressed with their known physically interacting partners or regulatory targets in ASD leukocytes. Accordingly, the XP-ASD network included a total of 316 out of 965 (36%) likely rASD genes.

The 965 rASD genes included both high-confidence rASD genes (e.g., recurrently mutated in individuals with ASD) and low-confidence rASD genes (some even found in siblings of individuals with ASD, who developed normally). We reasoned that if the XP-ASD network is truly relevant to the prenatal etiology of ASD, high-confidence rASD genes would be preferentially incorporated into the XP-ASD network. By following different analytical methods, other researchers have independently categorized rASD genes into high- and low-confidence^7, 14, 42^. Importantly, we found a reproducible enrichment of high-confidence rASD genes in the XP-ASD network (Fig. 3c). We also observed a significant enrichment for strong-candidate rASD genes with *de novo* protein truncating variants in the XP-ASD network (hypergeometric p-value <3.6×10^-6^). Further corroborating a possible regulatory role of rASD genes on the DE-ASD network, rASD genes in the XP-ASD network were significantly enriched for DNA-binding activity, compared to the remaining rASD genes (OR: 3.1; p-value <2.1×10^-12^; Fisher’s exact test; Fig. S7). Furthermore, the XP-ASD network was not enriched for rASD genes classified as low-confidence (p-value >0.24; SFARI categories 4-6). As negative controls, we constructed two other networks by including genes with likely deleterious and synonymous mutations in siblings of individuals with ASD, who developed normally^13^. Consistent with a possible role of the XP-ASD network in ASD, these negative control genes were not significantly associated with the DE genes (p-values >0.41; Fig. 3c). The preferential addition of high-confidence and regulatory rASD genes supports the relevance of the XP-ASD network for the pathobiology of ASD, and the likelihood that the high-confidence rASD genes are regulating the DE-ASD network.

### rASD genes tend to be repressors of genes in the DE-ASD network

To explore how rASD genes may regulate DE genes, we analyzed their interaction types (i.e., positive or negative correlations, alluding to activator or repressor activity). Comparative analysis of interactions between DE and rASD genes in the XP-ASD network indicated a significant enrichment of negative correlations between rASD and DE genes (OR: 1.79; p-value <3.1×10^-4^; Fisher’s exact test), suggesting a predominantly inhibitory role of rASD genes on the DE genes (Fig. 4a).

**Fig 4.**
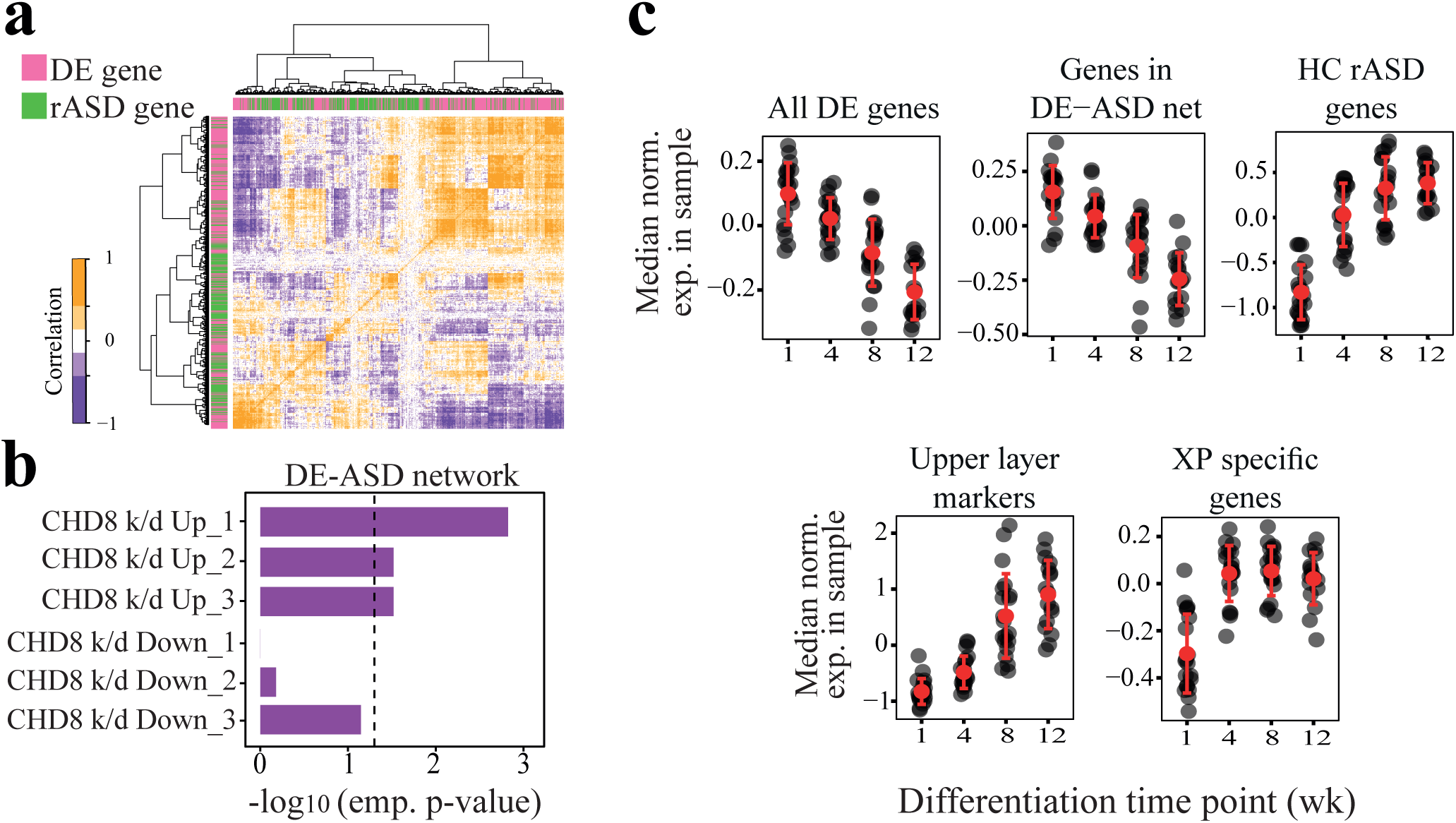
rASD genes potentially suppress the DE genes. a) Interactions between DE and rASD genes are enriched for negative Pearson’s correlation coefficients in the ASD leukocyte transcriptome (n=119 subjects; see Fig. S7 for more details). b) The DE-ASD network is significantly enriched for genes that are up-regulated following the knock-down of CHD8 (empirical tests). Data were extracted from three studies: Sugathan et al.^36^ (*CHD8* k/d_1), Gompers et al.^38^ (*CHD8* k/d_2), and Cotney et al.^37^ (*CHD8* k/d_3). See also Fig. S8. c) Expression patterns of DE-ASD genes were negatively correlated with rASD genes during *in vitro* differentiation of human primary neural precursor cells^43^ (n= 77 samples across time points; 3 fetal brain donors). In each panel, black circles represent the median expression of associated genes in a sample. Expression levels of each gene were normalized to have mean of zero and standard deviation of one across samples. While genes in the DE-ASD network are significantly down-regulated during neuron differentiation (p-value = 4.4 x 10^-6^; Wilcoxon-Mann-Whitney test), XP specific genes are significantly up-regulated (p-value = 1.2×10^-3^; Wilcoxon-Mann-Whitney test). The expression levels of CACNA1E, PRSS12, and CARTPT were considered as the markers of upper layer neurons (late stage of neural differentiation). See Fig. S7 for related details.

In line with a role of rASD genes as repressors, the DE-ASD network was enriched for genes that were up-regulated by the knock-down of *CHD8* in neural progenitor and stem cells, but not for genes that were down-regulated^36–38^ (Fig. 4b). Consistent with this, gene set enrichment analysis demonstrated an overall up-regulation of genes that are also up-regulated in knock-down experiments of the transcriptional repressor *CHD8* (p-value <0.039 across three different studies^36–38^), but not for those that are down-regulated. There was a similar trend towards up-regulation for the binding targets of the *FMR1* rASD gene in the ASD transcriptome^39^ (p-value: 0.078; GSEA).

To further test if rASD genes were predominantly repressors of genes in the DE-ASD network, we analyzed an independent transcriptome dataset from the differentiation of primary human neural progenitor cells obtained from fetal brains of three donors^43^. We found that expression of genes in the DE-ASD network exhibit a gradual down-regulation during neural progenitor differentiation (p-value 4.4×10^-6^; Fig. 4c). However, the genes unique to the XP-ASD network (i.e., rASD genes present in the XP-ASD network, but not DE-ASD network) showed an anti-correlated expression pattern with DE-ASD genes with peak expression at 12 weeks into differentiation (p-value 1.2×10^-3^; Fig. 4c). The results of this independent dataset provide further evidence of a potential inhibitory role of rASD genes on DE-ASD networks during human neuron differentiation.

### Signaling pathways are central to the leukocyte-based networks

We next identified key pathways involved in the XP-ASD and DE-ASD networks. Biological process enrichment analysis of the XP-ASD network demonstrated it is highly enriched for signaling pathways (Fig. 5a). Moreover, the DE-ASD network was highly enriched for PI3K/AKT, mTOR, and related pathways (Fig. 5b). To delineate mechanisms by which rASD genes could dysregulate DE genes, we compared enriched biological processes between DE and rASD genes in the XP-ASD network. DE genes were more enriched for cell proliferation-related processes, particularly PI3K/AKT and its downstream pathways such as mTOR, autophagy, viral translation, and FC receptor signaling (Fig. 5a-b). However, the rASD genes were more enriched for processes involved in neuron differentiation and maturation, including neurogenesis, dendrite development and synapse assembly (Fig. 5a).

**Fig 5.**
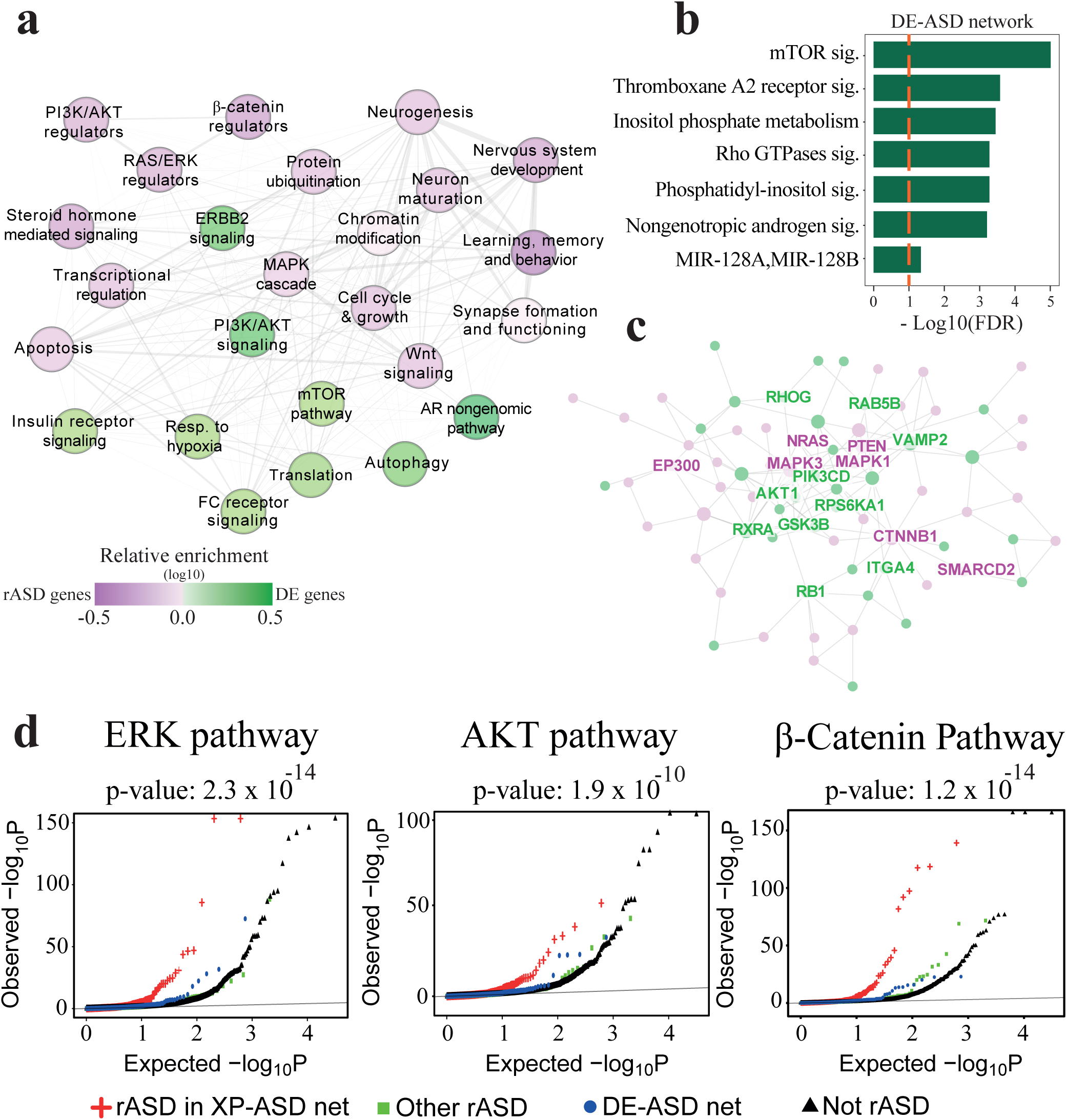
The architecture of the XP-ASD network implicates PI3K/AKT, RAS/ERK, WNT/β-catenin pathways as core dysregulated processes in ASD, regulated by rASD genes. a) Summary of enriched biological processes in the XP-ASD network. Each node represents a biological process that is significantly enriched in the XP-ASD network (two-sided Fisher’s exact test). Nodes that preferentially include rASD and DE genes are represented by purple and green colors, respectively. The interactions among terms represent the connection patterns of their cognate genes in the XP-ASD network with thicker interactions indicating more significant connections (hypergeometric test). Only connections with p-value <0.05 are shown. This illustration covers 86% of genes involved in the XP-ASD network. b) Significantly enriched processes in the DE-ASD network (Benjamini-Hochberg corrected FDR <0.1; hypergeometric test). These processes are also up-regulated in ASD leukocytes based on GSEA (n=119 ASD and 107 TD). c) The connected graph of hubs in the XP-ASD network. Green nodes represent hub genes in both XP-ASD and DE-ASD networks, while XP-ASD network-only hub genes are in purple. See Fig. S9 for the network with all gene labels. d) Significant enrichment of rASD genes in the XP-ASD network for the regulators of RAS/ERK, PI3K/AKT, WNT/β-catenin pathways. The x-axis indicates the p-value that gene mutations would dysregulate the corresponding signaling pathways. The background is composed of all genes that were assayed in Brockmann et al.^47^, excluding rASD and DE genes. The significance of enrichment of rASD genes in XP-ASD network for the regulators of signaling pathways were examined using Wilcoxon-Mann-Whitney test with background genes (illustrated in black) as control.

Our results suggest elevated co-expression activity of PI3K/AKT and its downstream pathways in ASD leukocytes (Fig. 5a-b). These processes are involved in brain development and growth during prenatal and early postnatal ages^3, 45, 46^ and focused studies on rASD genes have implicated them in ASD^3, 10, 44–46^. Further supporting the increased co-expression activity of the PI3K/AKT and its downstream pathways in our cohort of toddlers with ASD, gene set enrichment analysis demonstrated that the PI3K/AKT pathway and two of its main downstream processes (upregulation of mTOR pathway and upregulation of genes that are regulated by FOXO1) are also dysregulated in ASD leukocytes in directions that are consistent with the increased activity of the PI3K/AKT pathway.

We further investigated the DE-ASD and XP-ASD networks using an integrated hub analysis approach (Methods). In the DE-ASD network, hub genes included the key members of the PI3K/AKT pathway including *PIK3CD*, *AKT1* and *GSK3B* (Fig. 5c; Fig. S8). Genes that were only hubs in the XP-ASD network included regulators of neuronal proliferation and maturation, including regulatory members of the RAS/ERK (e.g., *NRAS*, *ERK2*, *ERK1*, *SHC1*), PI3K/AKT (e.g., *PTEN*, *PIK3R1*, *EP300*), and WNT/β-catenin (e.g., *CTNNB1*, *SMARCC2*, *CSNK1G2*) signaling pathways (Fig. 5c; Fig. S9). While PI3K/AKT (a hub in DE-ASD and XP-ASD networks) promotes proliferation and survival, many of the genes that are only hub in the XP-ASD network, including *NRAS*, *ERK 1/2*, and *PTEN,* can trigger differentiation of neural progenitor cells by mediating PI3K/AKT and its downstream pathways^3, 44^.

### rASD genes regulate DE-ASD genes through specific signaling pathways

We further explored if perturbation to the rASD genes lead to the perturbation of the DE-ASD network through changes in the RAS/ERK, PI3K/AKT, and WNT/β-catenin pathways. The activity of these three pathways is chiefly mediated through changes in phosphorylation of ERK, AKT, and β-catenin proteins. Therefore, to assess the regulatory influence of rASD genes on these signaling pathways, we leveraged available genome-wide mutational screening data wherein gene mutations were scored based on their effects on the phosphorylation state of ERK, AKT, and β-catenin proteins^47^. Consistent with the functional enrichment and hub analysis results, rASD genes in the XP-ASD network were significantly enriched for regulators of the RAS/ERK, PI3K/AKT, and WNT/β-catenin pathways (Fig. 5d; p-value <1.9×10^-10^). Specifically, regulators of these pathways (FDR <0.1) accounted for inclusion of 39% of rASD genes in the XP-ASD network. No significant enrichment for regulators of the RAS/ERK, PI3K/AKT, and WNT/β-catenin pathways was observed among rASD genes that were not included in the XP-ASD network (Fig. 5d). These results support the notion that rASD genes regulate the DE-ASD network through perturbation of the RAS/ERK, PI3K/AKT, and WNT/β-catenin pathways.

In summary, our XP-ASD network decomposition results suggest a modular regulatory structure for the XP-ASD network in which diverse rASD genes converge upon and dysregulate activity of the DE genes (Fig. 5a). Importantly, for a large percentage of rASD genes, the dysregulation flow to the DE genes is channeled through highly inter-connected signaling pathways including RAS/ERK, PI3K/AKT, and WNT/β-catenin.

### The DE-ASD network is over-active in neuron models of individuals with ASD and brain enlargement

Our results demonstrate increased gene co-expression in the DE-ASD network in leukocytes of toddlers with ASD selected from the general population. Furthermore, they implicate the DE-ASD network in the prenatal etiology of ASD by demonstrating its higher co-expression during fetal brain development, and its connection with high-confidence rASD genes. Also, our results suggest that the increased co-expression in the network is present in a large percentage of our ASD toddlers and is associated with the processes related to the neural proliferation and maturation.

To further validate these results, we examined if the DE-ASD network shows increased co-expression in hiPSC-derived neural progenitors and neurons from toddlers with ASD. Thus, we re-analyzed a previously published hiPSCs transcriptome data from 13 individuals with ASD and TD^28, 48^, which were differentiated into neural progenitor and neuron stages. The included subjects with ASD capture macrocephaly which is an important phenotype common in many subjects with ASD. Importantly, our analysis demonstrated that the DE-ASD network is more active in these neuron models of subjects with ASD (Fig. 6; Fig. S10). This result suggests the functional relevance of identified leukocyte molecular signatures to the abnormal brain development in ASD, particularly for individuals with brain enlargement.

**Fig 6.**
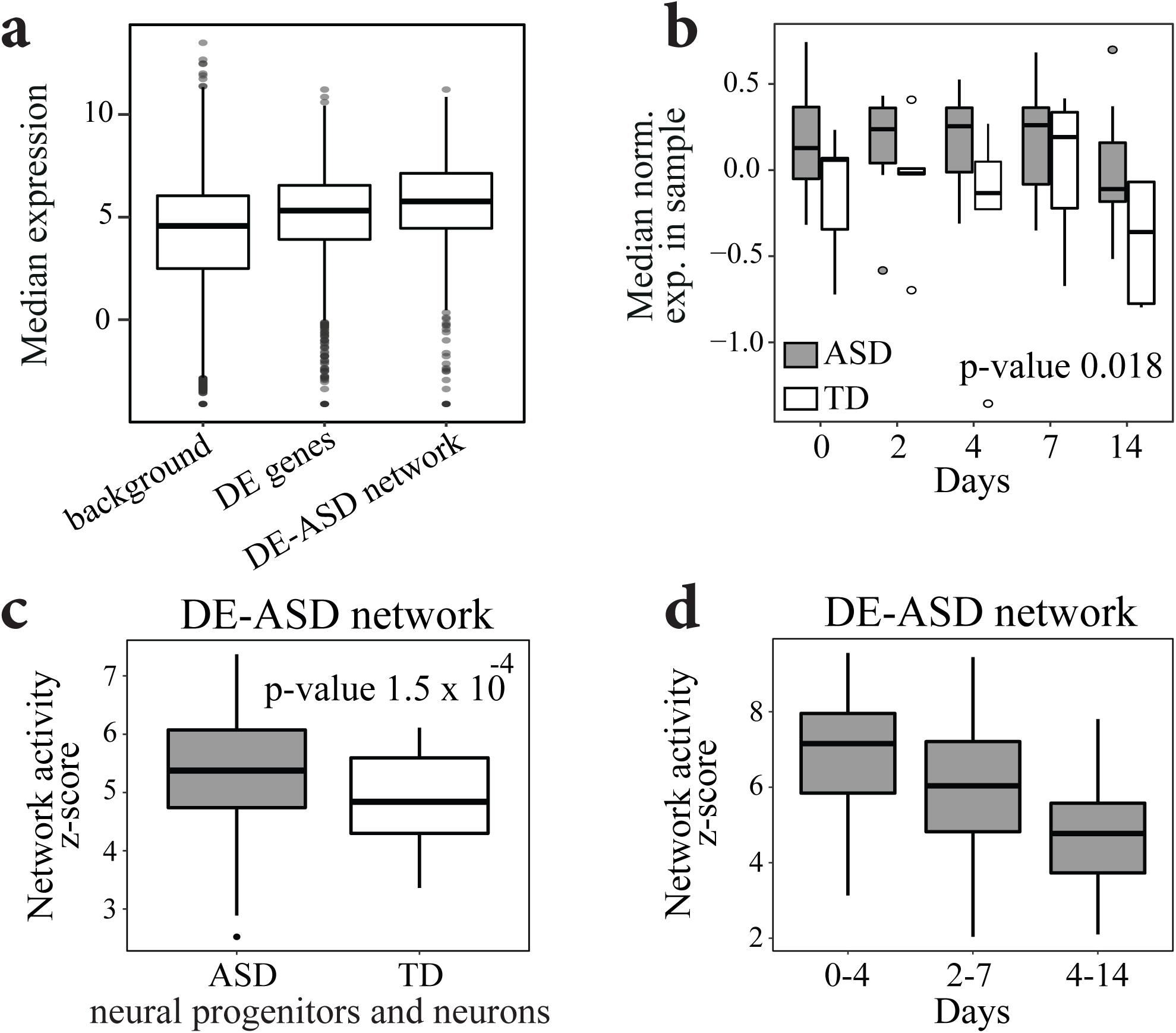
The DE-ASD network show increased gene co-expression in differentiating neurons of ASD. a) The genes in the DE-ASD network are highly expressed during neural differentiation of hiPSCs from ASD and TD cases^48^ (p-value 7.4 x 10^-25^; two-sided Wilcoxon-Mann-Whitney test). For each gene, its median normalized expression at neural progenitor and neuron stages was considered (n= 65 hiPSC-derived neural progenitor and neuron samples from 13 donors). Similar patterns were observed when analyzing each stage independently. b) The DE-ASD network shows higher expression level in hiPSC-derived neural progenitor and neuron stages of individuals with ASD. Expression data for each gene was normalized to have mean zero and variance of one. Boxplots represent the distribution of median expression levels genes involved in the DE-ASD network at each differentiation time point. A mixed linear regression with subjects as random effects was used to estimate the significance of the observed pattern (n=65 samples from 13 donors across 5 time points). This demonstrated the diagnosis group as a significant factor in expression level of genes involved in the DE-ASD network (p-value: 0.018; estimate: 0.31; std. err.: 0.13; t-statistics: 2.36; df: 57). c) The DE-ASD network shows higher co-expression activity in ASD derived neural progenitors and neurons. To estimate the co-expression strength of interacting gene pairs in DE-ASD network in neural progenitor and neurons of in each diagnosis group, iterating 100 times, we randomly selected 4 individuals from a diagnosis group and measured the co-expression strength of the DE-ASD network at neural progenitor differentiation time points of day 0, 2, 4, 7, and 14 (n=20 samples). The boxplots represent the distribution of z-transformed p-values of co-expression strength as measured by a two-sided Wilcoxon-Mann-Whitney test. d)The highest co-expression activity of the DE-ASD network in subjects with ASD coincides with the proliferation period (Day 0-to-4) of neural progenitors, and then its transcriptional activity gradually decreases. See Fig. S10 for more details. Boxplots represent the median (horizontal line), lower and upper quartile values (box), and the range of values (whisker), and outliers (dots).

### Network dysregulation is associated with ASD severity

We evaluated the potential role of the DE-ASD network activity on the development of the core clinical symptom of socialization deficits in toddlers with ASD. To this end, we first tested if the same pattern of gene co-expression dysregulation exists across individuals at different levels of ASD severity as measured by Autism Diagnostic Observation Schedule (ADOS) social affect severity score. We observed that the fold change patterns of DE genes are almost identical across different ASD severity levels (Fig. S11). The implicated RAS/ERK, PI3K/AKT, WNT/β-catenin pathways in our model are well known to have pleotropic roles during brain development, from neural proliferation and neurogenesis to neural migration and maturation. These signaling pathways and the associated developmental stages have been implicated in ASD^3^, suggesting the DE-ASD network is involved in various neurodevelopmental processes. At the mechanistic level, this suggests that the spectrum of autism could reflect the varying extent of dysregulation of the DE-ASD network, as it is composed of high-confidence physical and regulatory interactions. Hence, we examined whether the magnitude of the co-expression activity of the DE-ASD network correlated with clinical severity in toddlers with ASD. Indeed, we found that the extent of gene co-expression activity within the DE-ASD network was correlated with ADOS social affect deficit scores of toddlers with ASD (Fig. 7). To assess the significance of observed correlation patterns, we repeated the analysis with 10,000 permutations of the ADOS social affect scores in individuals with ASD. This analysis confirmed the significance of the observed correlations (inset boxplots in Fig. 7). Our results suggest the perturbation of the same network at different extents can potentially result in a spectrum of postnatal clinical severity levels in toddlers with ASD.

**Fig 7.**
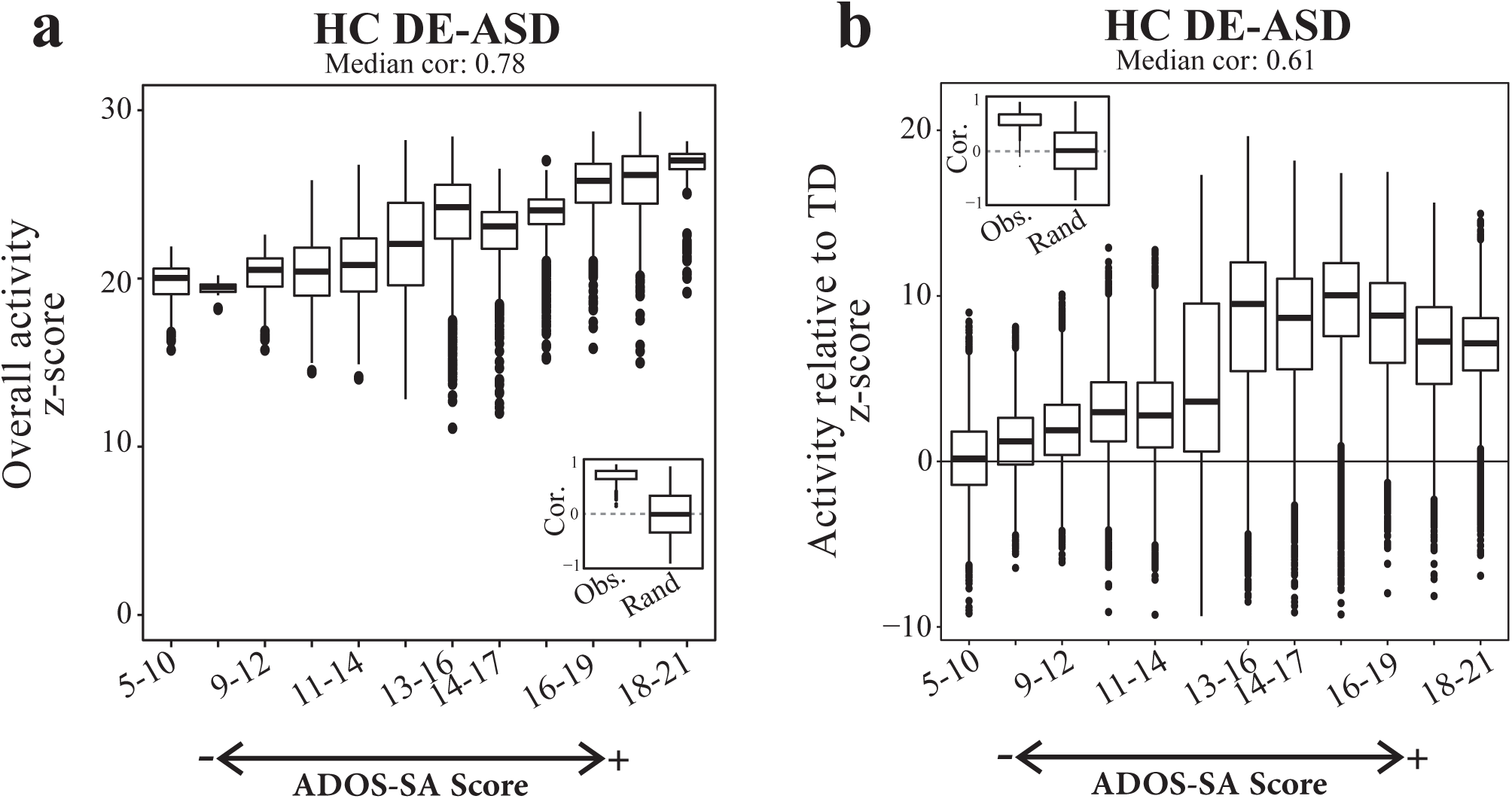
Co-expression magnitude of the DE-ASD network correlates with ASD severity. a) ASD toddlers were sorted by their ADOS social affect scores (ADOS-SA) with higher scores representing more severe cases. The network activity was measured in a running window on ADOS-SA scores. The overall activity of the DE-ASD network in a set of samples was measured by comparing the co-expression magnitude of interactions in the network with the background derived from the same set of samples (Methods). To ensure robustness of the results, we measured the co-expression activity of the DE-ASD network at each severity group by randomly selecting n=20 subjects with ASD from that severity level, iterating 1000 times. The left inset panel illustrates the distribution of observed correlation values of DE-ASD network with the ADOS-SA severity, and compares it with permuted data from10,000 random shuffling of ADOS-SA scores of subjects with ASD (two-sided p-value <10^-6^; permutation test; see Fig. S11). b) The relative co-expression magnitude of the DE-ASD networks compared to TD cases. The relative activity level was estimated by comparing the co-expression strength of interactions in the DE-ASD network between ASD and TD toddlers. For each severity group, n=20 ASD samples in that ADOS-SA range were randomly selected and compared to n=20 random TD samples, iterating 1000 times. Significance of the trend was evaluated by 10,000 permutations of the ADOS-SA scores in toddlers with ASD (two-sided p-value <10^-6^; permutation test; see Fig. S11-S12). Boxplots represent the median (horizontal line), lower and upper quartile values (box), and the range of values (whisker), and outliers (dots).

## Discussion

While ASD has a strong genetic basis, it remains elusive how rASD genes are connected to the molecular changes underlying the disorder at prenatal and early postnatal ages. We developed a systems-biology framework to identify perturbed transcriptional programs in leukocytes, and connect them with the rASD genes and early-age symptom severity. Specifically, we found a dysregulated gene network that shows elevated gene co-expression activity in leukocytes from toddlers with ASD. This core network was robustly associated with high-confidence rASD genes. Although recurrent, high confidence rASD gene mutations occur in a small percentage of the ASD population^5, 14^. The connection of the DE-ASD network (constructed with data from the general ASD pediatric population) with high-confidence rASD genes provides evidence of shared mechanisms underlying ASD in both individuals with highly penetrant rASD gene mutations and those with other etiologies (e.g., common variants). We further show that many rASD genes may regulate the DE-ASD core network through the RAS/ERK, PI3K/AKT, and WNT/β-catenin signaling pathways. This study confirms and substantially expands results from previous reports on blood transcriptome of subjects with ASD.

A key aspect of our signature is that it allows one to investigate the relationship of molecular perturbations with early-age ASD symptom severity. Indeed, we found that the magnitude of dysregulation of the DE-ASD network is correlated with deficits in ADOS social affect scores in male toddlers of 1-4 years old. Social and behavioral deficits are also suggested to be linked with the genetic variations in subjects with ASD^49, 50^; and previous studies have established the effect of the PI3K/AKT signaling pathway (central to the DE-ASD core network) on social behaviors in mouse models^45, 46^. Together, these observations suggest that the etiology of ASD converges on gene networks that correlate with ASD symptom severity. Moreover, our results reinforce the hypothesis that stronger dysregulation of this core network could lead to a higher ASD severity. The DE-ASD core network is enriched for pathways implicated in ASD, strongly associated with high-confidence rASD genes, and correlate with the ASD symptom severity. However, a direct causal relationship between the co-expression activity of the network and ASD remains to be established. Moreover, our co-expression activity measure is a summary score from the strongest signal in our dataset (i.e., differentially expressed genes) at a group level (i.e., severity level). Therefore, by design, it may not comprehensively capture the heterogeneity that could exist within ASD. Future work is needed to explore the causal relationship of the pathways in the DE-ASD network to ASD development, symptoms, and the potential existence of other dysregulation mechanisms in individuals with ASD.

Emerging models of complex traits suggest that gene mutations and epigenetic changes often propagate their effects through regulatory networks and converge on core pathways relevant to the trait^21, 30^. Our findings support the existence of an analogous architecture for ASD, wherein rASD genes with diverse biological roles converge and regulate core downstream pathways. Although the DE-ASD network did not significantly overlap with rASD genes, we found that it was significantly co-expressed with rASD genes in both leukocyte and brain. We also showed that the DE-ASD network genes are regulated by many rASD genes through direct transcriptional regulation or by modulating highly interconnected signaling pathways. We postulate that the DE-ASD network is a primary convergence point of ASD etiologies. This predicts that the spectrum of autism in such cases reflects degree and mechanism of the perturbation of the DE-ASD network. A detailed analysis of hiPSC-derived neurons from subjects with ASD and brain enlargement demonstrated the dysregulation of the DE-ASD network in these neuron models of ASD. Furthermore, clinical relevance is demonstrated by the high correlation we found between magnitude of dysregulation in the DE-ASD core network and ASD symptom severity in the toddlers.

The vast majority of rASD genes are not fully penetrant to the disorder^3, 8, 14^. Our analysis of the XP-ASD network sheds light on how rASD genes could potentially combine to result in ASD. Although some rASD genes could directly modulate the DE-ASD network at the transcriptional level, our results suggest that the regulatory consequence of many rASD genes on the DE-ASD network are channeled through the PI3K/AKT, RAS/ERK, WNT/β-catenin signaling pathways. The structural and functional interrogation of the XP-ASD network localized these pathways to its epicenter and demonstrated enrichment for processes downstream of these pathways among DE genes. Moreover, we found that high-confidence rASD genes are better connected to the DE-ASD core network, suggesting that the closeness and influence of genes on these signaling pathways is correlated with their effect size on the disorder. These results articulate that perturbation of the PI3K/AKT, RAS/ERK, WNT/β-catenin signaling pathways through gene regulatory networks may be an important etiological route for ASD that could be associated with the disorder severity level in a large fraction of the ASD population. Congruent with this hypothesis, cellular and animal models of ASD have demonstrated that high-confidence rASD genes are enriched in regulators of the RAS/ERK, PI3K/AKT, WNT/β-catenin signaling pathways^3, 10^. These signaling pathways are highly conserved and pleiotropic, impacting multiple prenatal and early postnatal neural development stages from proliferation/differentiation to synaptic and neural circuit development^3^. Such multi-functionalities could be the reason that we detected the signal in leukocytes of individuals with ASD.

It is necessary to analyze large subject cohorts from unbiased, general pediatric community settings to capture the heterogeneity that underlies ASD at early ages. This study presents the largest transcriptome analysis thus far from such settings. However, the analyzed dataset is still of a modest size, and as such our analysis focused on the strongest signal that best differentiates ASD and TD diagnosis groups (i.e., differentially expressed genes). Here we illustrate that the captured signal is informative about the transcriptional organization underlying ASD and shows promise in bridging the gap between genetic and clinical outcomes. Future studies with larger datasets are required to not only replicate these results, but also explore other long-standing questions in the field, such as the basis of sex bias that exists in ASD or the potential molecular mechanisms that differentiate high-functioning from low-functioning individuals.

Finally, current ASD diagnostics rely solely on behavioral phenotypes of toddlers, and such approaches are limited by age wherein ASD can be reliably diagnosed^3, 16^. Hence, an exciting direction is to expand the presented framework to systematically diagnose, classify and prognostically stratify subjects with ASD at earlier postnatal ages based on mRNA or other molecular markers. The concept of precision molecular medicine for ASD can only be actualized via approaches that illuminate the early-age molecular basis of ASD^3, 19^. Emerging evidence represent ASD as a progressive disorder that, at prenatal and early postnatal stages, involves a cascade of diverse varying molecular and cellular changes, such as those resulting from dysregulation of the pathways and networks described in this paper^3, 28, 29^. As such, it will be invaluable to develop molecular assays to assess infants and toddlers. The framework presented here could facilitate the development of such measures for ASD diagnosis and prognosis, by identifying specific molecular dysregulations that we show are observable in leukocytes of a large fraction of toddlers with ASD.

## Data availability

Leukocyte transcriptome data can be accessed from the NCBI Gene Expression Omnibus (GEO) database under the accession codes of GSE42133 and GSE111175. Microarray transcriptome data on the differentiation of primary human neural progenitor cells to neural cells were downloaded from the NCBI GEO accession GSE57595. Transcriptome data on hiPSC-derived neuron models of ASD and TD were downloaded from EMBL-EBI ArrayExpress with the accession code E-MTAB-6018. Human brain developmental transcriptome data were downloaded from BrainSpan.org.

## Accession codes

Gene Expression Omnibus database (GSE42133; GSE111175; GSE57595). EMBL-EBI ArrayExpress (E-MTAB-6018).

## Code availability

The R code for reproducing the analyses reported in this article is available at https://gitlab.com/LewisLabUCSD/ASD_Transcriptional_Organization.

## Acknowledgments

Authors would like to thank Dr. Lilia Iakoucheva for her helpful comments on this manuscript. This work was supported by NIMH R01-MH110558 (E.C., N.E.L.), NIMH R01-MH080134 (K.P.), NIMH R01-MH104446 (K.P.), NFAR grant (K.P.), NIMH P50-MH081755 (E.C.), Brain & Behavior Research Foundation NARSAD (T.P.), and generous funding from the Novo Nordisk Foundation through Center for Biosustainability at the Technical University of Denmark (NNF10CC1016517 N.E.L.).

## Author Contributions

V.H.G., T.P., K.P., E.C., and N.E.L. conceived the project and designed the experiments. T.P., S.N., S.M., and L.L. collected the samples, conducted transcriptome assays, and managed the data. V.H.G. and B.P.K. analyzed the data. V.H.G., T.P., E.C., and N.E.L. interpreted the results and wrote the manuscript. E.C. and N.E.L. supervised the project.

## Materials and Methods

### Participant recruitment and clinical evaluation

In this study, we performed transcriptomics analysis of 302 male toddlers with the age range of 1 to 4 years. The samples were divided into primary discovery and replication datasets, and were assayed by either microarray or RNA-Seq platforms. This included previously published transcriptome data (153 individuals)^19^ and new samples were collected using a similar methodology for participant recruitment (149 new cases). Research procedures were approved by the Institutional Review Board of the University of California, San Diego. Parents of subjects underwent Informed Consent Procedures with a psychologist or study coordinator at the time of their child’s enrollment.

About 70% of toddlers were recruited from the general population as young as 12 months using an early detection strategy called the 1-Year Well-Baby Check-Up Approach^51^. Using this approach, toddlers who failed a broadband screen, the CSBS IT Checklist^52^, at well-baby visits in the general pediatric community settings were referred to our Center for a comprehensive evaluation. The remaining subjects were obtained by general community referrals. All toddlers received a battery of standardized psychometric tests by highly experienced Ph.D. level psychologists including the Autism Diagnostic Observation Schedule (ADOS; Module T, 1 or 2), the Mullen Scales of Early Learning and the Vineland Adaptive Behavior Scales. Testing sessions routinely lasted 4 hours and occurred across 2 separate days. Toddlers younger than 36 months in age at the time of initial clinical evaluation were followed longitudinally approximately every 9 months until a final diagnosis was determined at age 2-4 years. For analysis purposes, toddlers (median age, 27 months) were categorized into two groups based on their *final* diagnosis assessment: 1) ASD: subjects with the diagnosis of ASD or ASD features; 2) TD: toddlers with typical developments.

ADOS scores at each toddler’s final visit were used for correlation analyses with DE-ASD network co-expression activity scores. All but 4 toddlers were tracked and diagnosed using the appropriate module of the ADOS (i.e., ADOS Module-Toddler, Module-1, or Module-2) between the ages of 24-49 months, an age where the diagnosis of ASD is relatively stable^16^; the remaining 4 toddlers had their final diagnostic evaluation between the ages of 18 to 24 months.

### Blood sample collection

Blood samples were usually taken at the end of the clinical evaluation sessions. To monitor health status, the temperature of each toddler was monitored using an ear digital thermometer immediately preceding the blood draw. The blood draw was scheduled for a different day when the temperature was higher than 99 Fahrenheit. Moreover, blood draw was not taken if a toddler had some illness (e.g., cold or flu), as observed by us or stated by parents. We collected four to six milliliters of blood into ethylenediaminetetraacetic-coated tubes from all toddlers. Blood leukocytes were captured and stabilized by LeukoLOCK filters (Ambion) and were immediately placed in a −20°C freezer. Total RNA was extracted following standard procedures and manufacturer’s instructions (Ambion).

### Data processing and differential gene expression analysis of the primary dataset

The primary discovery dataset composed of 275 samples from 240 male toddlers with the diagnosis of ASD and TD from the general population. Gene expressions were assayed using Illumina HT-12 platform. All arrays were scanned with the Illumina BeadArray Reader and read into Illumina GenomeStudio software (version 1.1.1). Raw Illumina probe intensities were converted to expression values using the lumi package^53^. We employed a three-step procedure to filter for probes with reliable expression levels. First, we only retained probes that met the detection p-value <0.05 cut-off threshold in at least 3 samples. Second, we required the probes to have expression levels above 95^th^ percentile of negative probes in at least 50% of samples. The probes with detection p-value >0.1 across all samples were selected as negative probes and their expression levels were pooled together to estimate the 95^th^ percentile expression level. Third, for genes represented by multiple probes, we considered the probe with highest mean expression level across our dataset, after quantile normalization of the data. These criteria led to the selection of 14,854 protein coding genes as expressed in our leukocyte transcriptome data, which is similar to the previously reported estimate of 14,555 protein coding genes (chosen based on unique Entrez IDs) for whole blood by GTEx consortium^54^. To ensure results are not affected by the variations in the procedure of selecting expressed genes, we replicated all of our analyses (redoing DE analysis and re-constructing HC DE and XP networks) by choosing 13,032 protein coding genes as expressed (Fig. S13).

Quality control analysis was performed on normalized gene expression data to identify and remove 22 outlier samples from the dataset. Samples were marked as outlier if they showed low signal intensity in the microarray (average signal of two standard deviations lower than the overall mean), deviant pairwise correlations, deviant cumulative distributions, deviant multidimensional scaling plots, or poor hierarchical clustering, as described elsewhere^18^. After removing low quality samples, the primary dataset had 253 samples from 226 male toddlers including 27 technical replicates. High reproducibility was observed across technical replicates (mean Spearman correlation of 0.917 and median of 0.925). We randomly removed one of each of two technical replicates from the dataset.

The limma package^55^ was then applied on quantile normalized data for differential expression analysis in which moderated t-statistics was calculate by robust empirical Bayes methods^56^. Sample batch was used as a categorical covariate (total of two batches; both Illumina HT-12 platforms). Exploration graphs indicated that linear modeling of batch covariate was effective at removing its influence on expression values (Fig. S14). MA-plots of the primary dataset did not show existence of bias in the fold change estimates (Fig. S1). DE analysis identified 1236 differentially expressed genes with Benjamini-Hochberg FDR <0.05.

### Reproducibility assessment using additional microarray and RNA-Seq datasets

We performed six analyses to confirm that fold change patterns of DE genes in the primary dataset are robust to alterations in the analysis pipeline, are not affected by the batches, and are present in the vast majority of samples. First, we included additional co-variates in our regression models. Second, the discovery dataset (253 high quality samples from 226 male toddlers) is an expanded version of a dataset that we analyzed and reported in our previous study using a different approach^1^. Therefore, we compared the fold changes from non-overlapping subjects from these two studies. Third, we performed Jack-knife resampling to confirm that the observed fold changes were not driven by a small subset of subjects, but rather shared by the vast majority of samples (Fig S1). We performed a similar analysis on the network activity levels and found that the higher co-transcriptional activity of DE-ASD networks are not driven by a small number of subjects (Fig S11). Fourth, we performed additional microarray transcriptome analyses to confirm that our results are replicable at technical and biological levels. We conducted transcriptome analysis on a second dataset composed of 56 randomly selected male toddlers from the primary dataset (35 ASD and 21 TD). We also analyzed a third microarray dataset composed of 48 male toddlers with 24 independent, non-overlapping toddlers with ASD, while 21 out of 24 TD cases overlapped with the primary dataset. These two datasets were assayed concurrently, but at a different time than the primary dataset. Moreover, in contrast to the primary dataset, the second and third datasets were assayed by Illumina WG-6 Chips. The pre-processing and downstream analysis of the second and third microarray datasets were conducted separately using the same approaches as the primary dataset.

Fifth, to further assess the reproducibility of the results across experimental platforms, we performed RNA-Seq experiments on 56 samples from an independent cohort of 12 (19 samples) TD and 23 (37 samples) male toddlers with ASD. None of these subjects overlapped with those in the microarray datasets. This allowed us to ensure our results are not subject nor platform (i.e., microarray vs. RNA-Seq) specific. RNA-Seq libraries were sequenced at the UCSD IGM genomics core on a HiSeq 4000. We processed the raw RNA-Seq data with our pipeline that starts with quality control with FastQC^57^. Low quality bases and adapters were removed using trimmomatic^58^. Reads were aligned to the genome using STAR^59^. STAR results were processed using Samtools^60^, and transcript quantification is done with HTseq-count^61^. Subsequently, low expressed genes were removed and data were log count per million (cpm) normalized (with prior read count of 1) using limma^55^. We performed SVA analysis^62^ on the normalized expression data and included the first surrogate variable as covariate to account for potential hidden confounding variables. Differential expression analysis was performed using the limma package with subjects modeled as random effects (Fig S4).

Finally, we explored the potential impact of variations in cell type composition^1^ and other hidden confounding variables^62^ on the results. As illustrated in Fig. S2, our results indicated that there is no statistically significant difference in the cell type composition between ASD and TD samples.

### ASD risk genes

ASD risk genes were extracted from the SFARI database^42^ on Dec. 7, 2016. We also included the reported risk genes from a recent meta-analysis, containing genes mutated in individuals with ASD but not present in Exome Aggregation Consortium database (ExAC)^14^. Together, these two resources provided 965 likely rASD genes that were used for the construction of the XP-ASD networks. Previously published genes with likely gene damaging and synonymous mutations in siblings of subjects with ASD, who developed normally were retrieved from Iossifov et al.^13^.

ASD high confidence risk genes were extracted from the SFARI database (genes with confidence levels of 1 and 2), Kosmicki et al.^14^ (recurrent gene mutations in individuals with ASD, but not present in the ExAC database), Sanders et al.^15^, and Chang et al.^7^. Strong evidence genes with *de novo* protein truncating variants in subjects with ASD were extracted from Kosmicki et al.^14^ and included rASD genes that were not in the ExAC database and have a probability of loss-of-function intolerance (pLI) score of above 0.9. Gene names in these datasets were converted to Entrez IDs using DAVID tools^63^.

To assess the overlap of DE-ASD networks with rASD genes, we considered our list of all rASD genes (965 genes), different lists of high confidence rASD genes (varying in size and composition) and their combinations, including all SFARI rASD genes, SFARI gene levels 1-to-3, SFARI gene levels 1 and 2, strong evidence rASD genes from Kosmicki et al.^14^, and strong evidence rASD genes from Sanders et al.^15^.

rASD genes with potential gene regulatory role were identified based on the gene ontology annotations. We considered an rASD as a potential gene regulator if it was annotated with either DNA-binding transcription factor activity (GO:0003700), DNA binding (GO:0003677), or DNA-templated regulation of transcription (GO:0006355).

### Construction of context specific networks

We first regressed out the interfering co-variate (i.e., batch group) from the quantile normalized expression values of the primary dataset (see the Data processing section). The Context Likelihood of Relatedness (CLR) algorithm^64^ was next applied on the batch corrected transcriptome data from ASD and TD diagnosis groups separately to construct two co-expression networks (technical replicates were randomly removed from the dataset prior to construction of the networks). The CLR algorithm employs a two-step procedure to infer significantly co-expressed gene pairs. First, it estimates the distribution of similarity scores for each gene based on the similarity that the gene shows with all other genes in the dataset using a mutual information metric. Second, it estimates the significance of the observed similarity score for each gene pair by testing how likely it is to have such a similarity score given the co-expression similarity score distributions of the two genes from the first step. The separate application of the CLR algorithm on ASD and TD samples provided global (i.e., all expressed genes) gene-gene co-expression similarity matrices for each diagnosis group. DE and expanded DE-and-rASD (XP) networks were next constructed from CLR-derived ASD and TD similarity matrices as detailed below.

To ensure the robustness of the results, we constructed three variants of the DE networks for each diagnosis group (i.e., ASD and TD; total of six networks). These networks varied in the number of nodes and edges, providing a tradeoff between sensitivity (number of false negative interactions) and specificity (number of false positive interactions) in our downstream analysis. Unless otherwise noted, we reported results that were reproducible in all three networks. The three networks include the high confidence network (HC; including strong evidence physical and regulatory interactions), the functional network (including interactions between previously known functionally related genes), and the full co-expression network. The full co-expression network is solely based on co-expression patterns of DE genes (i.e., all significantly co-expressed DE gene pairs with FDR <0.05 as judged by the CLR algorithm). To construct the HC and functional networks, we first retrieved the static HC and functional networks of the detected protein-coding DE genes from databases. The static HC network was obtained from the Pathway Commons database^65^ and was updated to include interactions from the most recent Reactome^66^ and BioGrid^67^ databases. The static functional network was extracted from the GeneMania webserver^68^ and included interactions supported by co-expression, protein-protein interactions, genetic interactions, co-localization, shared protein domains, and other predictions^68^. The backbone, static network of all DE-ASD and DE-TD networks composed of at least 96% DE genes. Static HC and functional networks were made context specific by retaining those database-derived interactions that were significantly co-expressed in the diagnosis group (The static backbone networks were shared between the DE-ASD and DE-TD networks). All figures in the main text are based on HC DE-ASD and DE-TD networks, and the results of functional and full co-expression networks are represented in the supplement.

By design, the HC network is smaller, more accurate, but potentially more biased as it includes genes that are more actively studied than those in the functional network. Both networks are smaller than the full co-expression network. Therefore, on average, the functional DE-ASD and DE-TD networks had 15x more interactions and 2.3x more genes than their HC counterparts. Similarly, the full DE-ASD and DE-TD networks had 6.4x more interactions and 1.05x more genes than their functional counterparts.

The XP-ASD networks were constructed using a similar approach, but from the union of protein-coding DE genes and 965 rASD genes. Our list of 965 rASD genes included genes that are ranked either as high confidence (supported with multiple studies or direct experimentation) or low confidence (some even have been found in healthy siblings of individuals with ASD). To assess the relevance of XP-ASD networks to the pathobiology of ASD, we also examined the association of XP-ASD networks with genes mutated in siblings of subjects with ASD, who developed normally. For this, we constructed two other variants of the XP-ASD networks by adding genes with likely gene damaging mutations (Siblings-LGD) and Synonymous (Siblings-Syn) mutations in our list of DE and rASD genes, separately. We next tested if these two variants of XP-ASD networks preferentially incorporated mutated genes in siblings of individuals with ASD, who developed normally. As the sole purpose of these two network variants were to test the relevance of the main XP-ASD network, they were not needed for follow up analyses. Similar to DE networks, the main figures represent results based on the HC XP-ASD network and the results for the functional and full XP-ASD networks are included in the supplement.

### Network and module overlap analysis

Unless otherwise noted, we used permutation tests to assess the significance of overlap between pairs of networks or modules. The background gene list for DE and XP networks were all protein coding genes that were expressed in our microarray experiments (see the gene expression preprocessing section for more details). DE genes did not show bias in terms of gene mutation rates (p-value=0.36; Wilcoxon-Mann-Whitney test) and length (p-value: 0.45; two-sided t-test). We extracted the gene mutation rates from a previous study^5^.

Empirical permutation tests were conducted by 10,000 random draws from background gene lists and measuring the overlaps. The actual overlap was then compared to the overlap distribution of random draws and an empirical p-value was estimated. In cases where the estimated empirical p-value was zero based on 10,000 permutation tests, we performed 90,000 additional random draws to obtain a more accurate estimation. If the estimated empirical p-value was still zero, a theoretical, hypergeometric-based p-value (non-zero) was considered. Multiple testing was corrected by the Benjamini-Hochberg procedure and FDR <0.1 was considered as significant, unless otherwise noted. By design, our functional and full DE and XP networks are highly sensitive and therefore include more than 90% of queried genes. Since we required replicable significant overlap of gene sets across our networks, this feature renders the overlap analysis robust to potential biases due to the network topology.

To identify genes that potentially regulate DE-ASD networks, we examined the overlap of DE-ASD networks with identified targets of human transcription factors as part of ENCODE^40^ and the curated Chea2016 database^41^. Overall, targets of 285 unique human transcription factors are assayed in the ENCODE and Chea2016 resources, and from these, 20 are currently annotated as high-confidence or suggestive evidence rASD genes by the SFARI database (SFARI categories 1 to 3). We performed overlap analysis between targets of transcription factors and each of the three DE-ASD networks separately using the hypergeometric test through the EnrichR portal^72^. Some of the transcription factors were assayed multiple times, providing partially different sets of target genes for these transcription factors. For such transcription factors, we had multiple p-values from the overlap analysis. Therefore, we used Fisher’s method to combine the enrichment p-values across assays related to a given transcription factor during the analysis of each DE-ASD network. Next, p-values were corrected using the Benjamini-Hochberg procedure. Only transcription factors whose targets were significantly enriched in all three DE-ASD networks were considered as significantly overlapping (FDR <0.1) with the DE-ASD networks. This resulted in the identification of 97 unique transcription factors whose targets are significantly enriched in all three DE-ASD networks. From these 97, 11 transcription factors are currently annotated as high confidence or suggestive evidence rASD genes. We assessed whether rASD genes are significantly enriched among the 97 transcription factors using a Fisher’s exact test.

### Hub analysis

The hub analysis of DE-ASD and XP-ASD networks were conducted by an integrated analysis of high-confidence (HC) and functional networks. By design, HC and functional networks each have their own advantages. Interactions in HC networks are presumably more accurate but potentially biased towards specific genes that are better studied. In contrast, hubs in functional networks are less susceptible to bias in knowledge on the interactome, but more prone to false positive interactions. Thus, we aimed to combine the information provided by the two networks to get a more accurate picture of hub genes. We first counted the number of interactions that each gene has in either of HC or functional networks. For the genes that were present in only one of the two networks, the interaction count of zero was considered for the other network. Then the p-value of hubness for each gene in a network (with the null hypothesis that the gene is not a hub) was determined by calculating the empirical probability of identifying a gene with the same number of interactions or higher in the network. Next, the hubness p-value score of each gene in HC and functional networks were combined together using Fisher’s method:

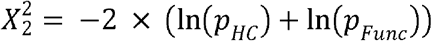

Where p refers to the empirical p-value of hubness for a gene in the HC and functional networks. [inine] is the chi-squared score with two degrees of freedom. The top 5% and 7% genes with highest [inline] scores were considered as hub in DE-ASD and XP-ASD networks, respectively.

### Functional characterization of DE-ASD networks

We set two criteria to identify biological processes that are differentially expressed between ASD and TD diagnosis groups and are enriched in the DE-ASD networks. First, we required the biological process to significantly change between ASD and TD transcriptome samples based on GSEA^69, 70^. Second, we required the biological process to be significantly enriched in the DE-ASD networks.

GSEA identified multiple gene sets that were significantly upregulated in subjects with ASD (FDR <0.12), using the R version of the GSEA package and the msigdb.v5.1 database (downloaded on Oct. 20, 2016)^69, 70^. Significantly enriched processes in the DE-ASD networks were identified by examining the overlap of GSEA-identified significantly altered gene sets with the DE-ASD networks based on empirical permutation tests, and p-values were corrected for multiple testing using the Benjamini-Hochberg procedure. We excluded gene sets associated with specific reference datasets in MSigDB since their generalizability to our dataset has not been established.

### Biological enrichment analysis of XP-ASD networks

Significantly enriched Gene Ontology biological processes (GO-BP) were identified by Fisher’s exact test on terms with the 10-2000 annotated genes. The terms with Benjamini-Hochberg estimated FDR <0.1 were deemed as significant. The enriched terms were next clustered based on the GO-BP tree, extracted from the Amigo database using RamiGO package in R^71^. The general terms with more than 1000 annotated genes that spanned two or more clusters were removed.

### Brain developmental gene expression data

Normalized RNA-Seq transcriptome data during human neurodevelopmental time periods were downloaded from the BrainSpan database on Dec. 20, 2016^34, 35^. To calculate correlations, normalized RPKM gene expression values were log2(x+1) transformed.

### Neural progenitor differentiation data

Microarray transcriptome data from the differentiation of primary human neural progenitor cells to neural cells^43^ were downloaded from the NCBI GEO database (GSE57595). The data were already quantile normalized and ComBat batch-corrected^73^. For genes with multiple probes, we retained the probe with the highest mean expression value.

To observe the transcriptome response of XP-ASD networks during neuron differentiation, we correlated the gene expression patterns with the developmental time points, considering the differentiation time as an ordinal variable.

### human induced pluripotent stem cells (hiPSC) data

We obtained hiPSC data^28^ from subjects with ASD and TD controls from EMBL-EBI ArrayExpress (E-MTAB-6018). Gene expression counts were normalized with the TMM method^74^ and filtered to exclude low-expressed genes (genes with count per million greater than 1 were retained). To calculate the correlations, normalized RNA-Seq gene expression values were log2(x+1) transformed. The subjects from this hiPSC study come from our center. However, none of the subjects overlap with those included in the transcriptome datasets in this study. Moreover, the hiPSC cohort includes only 8 subjects with ASD and macrocephaly, while our primary (i.e., discovery) leukocyte transcriptome is from 119 toddlers with ASD selected from general pediatric community and were not filtered based on their brain size. Moreover, the subjects participating in the two studies did not have the same age range and hiPSC cohort is composed of subjects with mean and median age of 167 and 193 months, respectively (toddlers in our dataset are between 12 to 48 months old). On the sample collection, our transcriptome data are from leukocytes of subjects with ASD, while the hiPSC transcriptome is based on the reprogrammed fibroblast cells.

### Regulatory effect of gene mutations on phosphorylation state of key signaling proteins

Data were extracted from a genome-wide mutational study that monitored the regulatory effect of gene mutations on phosphorylation status of 10 core genes of different signaling pathways and processes^17^. Genes whose mutations affected the phosphorylation status of the core signaling genes with FDR <0.1 were considered as the regulators of the cognate signaling pathway. We observed that 89 out of 316 rASD genes included in the HC DE-ASD network are regulators of at least one signaling pathway among RAS/ERK (as measured by phosphorylation status of ERKs), PI3K/AKT (as measured by phosphorylation status of AKT1), and WNT/β-catenin (as measured by phosphorylation status of β-catenin) (Fisher’s exact test p-value <1.7 x 10^-22^; see Fig 5d for more details). Moreover, 34 additional genes were connected to the XP-ASD network only through these 89 regulator genes. Therefore, as our HC networks are composed of high confidence interactions, in total 39% of rASD genes in the HC XP-ASD network are related to these three signaling pathways in the high confidence XP-ASD network. In general, signaling pathways are highly inter-connected in the cell^17^. Hence, genes could affect the activity of different pathways concurrently. To ensure that the genes included in the XP-ASD networks are specific to the three signaling pathways of RAS/ERK, PI3K/AKT, and WNT/β-catenin, we reproduced the results after removing the genes that regulate at least two signaling genes other than ERKs, AKT1, and β-catenin (data not shown). To observe the specificity of these results, as negative control, we confirmed that the rASD genes in the XP-ASD network are not significantly enriched for the regulators of LAMP1 gene after excluding those genes that regulate at least two of three signaling pathways of RAS/ERK, PI3K/AKT, and WNT/β-catenin.

### Measuring the co-expression activity of DE-ASD networks

We measured the co-expression strength of interacting genes in DE-ASD networks based on an unsigned Pearson’s correlation coefficient metric. To estimate the significance of the network activity in a set of samples, we compared the co-expression distribution of gene pairs in the network to a background distribution of co-expression values using the Wilcoxon-Mann-Whitney test in the R coin package. The network activity level was defined as z-transformed p-values of this comparison. Significant scores imply that at least some interacting gene pairs are co-expressed significantly higher than chance and hence parts of the network is potentially active. The background distribution was obtained by selecting genes with mean expression values closest to those involved in the relevant network. For each gene in the DE-ASD network, 10 non-overlapping genes with closest mean expression were selected. For example, we selected 3920 unique genes as the background for the HC DE-ASD network that has 392 genes. The unsigned correlations among these genes constituted the background distribution.

Measuring the co-expression activity of DE-ASD network during the neurodevelopmental period: To measure the co-expression activity of the DE-networks during brain neurodevelopmental periods from the BrainSpan RNA-Seq transcriptome data, we grouped samples from every 5 consequent time periods, starting from 8 post conception weeks and ending with 11 years old. The groups did not overlap in timespan.

Measuring the co-expression activity of DE-ASD network in neuron models of ASD: To measure DE-ASD network co-expression activity in hiPSC-derived neurons of ASD and TD cases, we analyzed the largest available transcriptome dataset including 8 ASD and 5 TD donors^6^. In this dataset, hiPSC-derived neural progenitor cells from each donor is differentiated to neurons and transcripts are measured during differentiation at 0, 2, 4, 7, and 14 days into the differentiation. To measure the DE-ASD network activity during the differentiation of neural progenitors to neuronal stages, iterating 100 times, we randomly selected 4 donors within each diagnosis group and measured the co-expression magnitude of the DE-ASD networks across the differentiation time points (n=20 samples).

Correlation of DE-ASD co-transcriptional activity with ADOS social affect scores: To map the co-expression activity of the DE-ASD networks on toddlers’ ADOS social affect (ADOS-SA) scores, we only considered ASD samples as DE-ASD networks were constructed based on genes that are differentially expressed between ASD and TD. toddlers with ASD were grouped based on a moving window on ADOS-SA scores with the width of 4 and a step size of 1. The number of toddlers with scores of 5 and 6 were relatively few compared to other categories. Therefore, the first window was from ADOS-SA score 5 to 10 (window size of 6). Moreover, to avoid biases due to number of samples in each window, the network activities were measured based on randomly selected sets of 20 samples from each window, iterating 1000 times. The correlation of ADOS-SA scores with the observed network activity was measured by considering the windows as ordinal values. To assess statistical significance, we shuffled the ADOS-SA scores of toddlers with ASD 10,000 times and re-calculated the network activity for each permutation (with no internal iterations).

There are some objective differences in measuring network activity during normal brain development versus the correlation of the leukocyte network activity with ADOS-SA scores. While in brain transcriptome data we hypothesized the DE-ASD networks show greater co-expression than background, we already knew that these networks are significantly co-expressed in toddlers with ASD and thus sought to test if their change in co-expression activity is dependent on ADOS-SA scores. Hence to map the relative activity of the DE-ASD networks in leukocytes of toddlers with ASD, in a second analysis, we based the background co-expression on the same network in the TD toddlers (instead of random genes from the same samples). The distribution of co-expression scores in each ADOS-SA score window was compared to the co-expression distribution (Wilcoxon-Mann-Whitney test) of the same network after randomly selecting the same number of samples among the TD toddlers (n=20 ASD samples and 20 TD samples at each iteration). We repeated the procedure 1000 times each with a distinct ASD and TD sample combination for all three context-specific DE-ASD networks to get the range of the network activity at each window. To assess the significance of observed distribution, we performed 10,000 permutations of ADOS-SA scores of toddlers with ASD (with no internal iterations at each severity level).

### Statistics and reproducibility

Almost all statistical analyses were conducted in the R programing environment (version 3.5.0). For microarray data, raw Illumina probe intensities were converted to expression values using the lumi package^53^. We filtered out probes that were not expressed from the dataset. Through quality control assessments, we identified and removed 22 outlier samples from the microarray dataset. Data were next quantile normalized and differentially expression genes were identified using limma package^55^ with the experimental batch included as a covariate in the regression model. Genes with FDR <0.05 were deemed as differentially expressed. Surrogate variable analysis did not support presence of other co-variates in the data^62^. Cibersort was used to examine potential impact of cell types on the differential expression patterns^75^. Technical replicates were used to assess the quality of samples and then were excluded from differential expression analysis and the follow up analyses (e.g., co-expression network construction). RNA-Seq data were mapped and quantified using STAR^59^ and HTSeq^61^, respectively. Quality of RNA-Seq samples were examined using FastQC^57^. Surrogate variable analysis was performed to identify and remove a covariate from RNA-Seq data^62^. Pearson’s correlation coefficient was used for the comparison of fold changes across datasets. We regressed out the covariate (i.e., the experimental batch) before calculating the co-expression. Significantly co-expressed genes were identified using the CLR package in MATLAB^64^, and interactions with co-expression FDR <0.05 were considered as significant. For network co-expression activity, we used unsigned Pearson’s correlation coefficient to measure the co-expression magnitude of interactions. The co-expression magnitudes of interactions of two networks were compared using two-sided Wilcoxon-Mann-Whitney test. When comparing co-expression magnitudes in two different datasets, to ascertain that the number of samples do not influence the measurements, a balanced number of samples were selected randomly. In most cases we used permutation tests to empirically examine the significance of an observed overlap between two gene sets. In cases that required a large number of tests, to increase speed, we used either hypergeometric or fisher’s exact tests. Fisher’s exact test was used to examine the overlap of the constructed networks with Gene Ontology-biological process (GO-BP) terms. We used the RamiGO package^71^ to cluster significantly enriched GO-BP terms that are similar and overlapping in their gene content. If appropriate, all p-values were corrected for multiple testing. The EnrichR portal^72^ was used to systematically examine the enrichment of the DE-ASD networks for the regulatory targets of human transcription factors. Fisher’s method was used to combine p-values from multiple assays on the same transcription factor. When applicable, we specified the sample sizes (*n)* within the figure legend or table description. Non-parametric tests (e.g., Wilcoxon-Mann-Whitney and permutation tests) were used to avoid strong assumptions about the distribution of data in our statistical analyses. No statistical tests were used to predetermine sample sizes, but our sample sizes were larger than those reported in previous publications^18, 19, 25^. No randomization was performed in our cohort assignment. Data collection and analysis were not performed blind to the conditions of the experiments.

