## Supplementary Figures for "A perturbed gene network containing PI3K/AKT, RAS/ERK, WNT/β-catenin pathways in leukocytes is linked to ASD genetics and symptom severity"

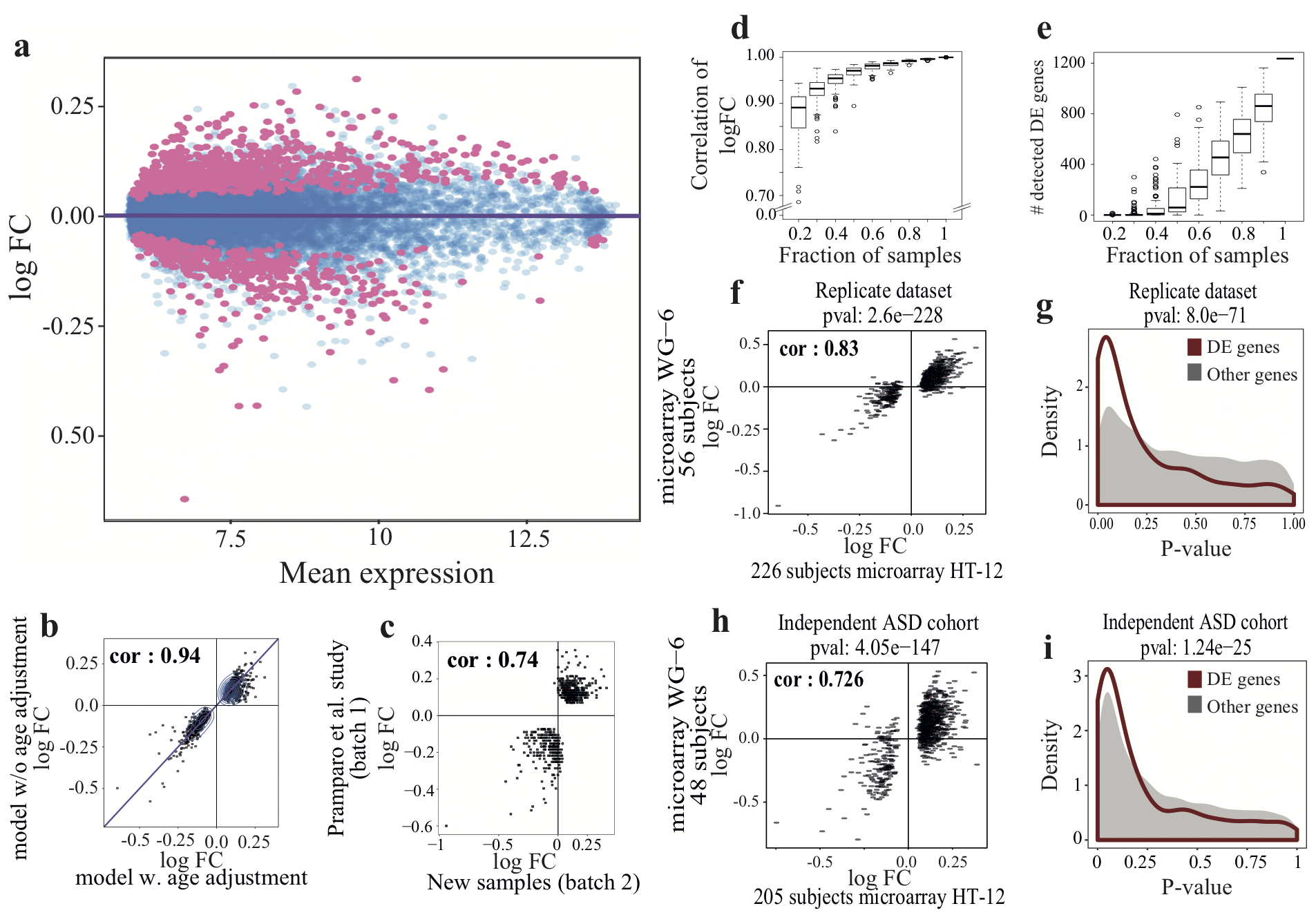

Supplementary Figure 1

Robustness analysis of observed DE patterns.

a) MA-plot of the primary dataset (n=119 ASD and 107 TD subjects). The purple line indicates the regression line between mean and fold change of all genes expressed in the dataset. As demonstrated, the mean expression and the fold changes are not correlated in overall. However, compared to all expressed genes, differentially expressed genes exhibit an up-regulation pattern (p-value <2.6x10^-63^; two-sided Wilcoxon-Mann-Whitney test). b) To observe the effect of the data processing approach on fold change patterns, the covariates of the linear regression model were changed in the limma package. As shown, similar fold change patterns were observed with or without inclusion of age as a covariate with a Pearson’s correlation coefficient of 0.94 (n= 119 ASD and 107 TD subjects). c) Our primary transcriptome dataset was composed of 226 subjects analyzed in two different batches including one batch of 128 samples (n= 84 ASD and 44 TD subjects) that was reported by Pramparo et al. previously and a second batch of 98 samples (n= 35 ASD and 63 TD subjects) that included new samples (samples non-overlapping between the batches; technical replicates were randomly removed). To assess whether the batch effect could be effectively handled, we compared the fold change patterns of DE genes between these two batches. For a more conservative analysis, we took the fold change of DE genes from the previously published batch from Pramparo et al^1^ study. These fold changes were calculated using a different analysis pipeline than the current study. We next compared those fold changes with our limma-based analysis of 98 new non-overlapping samples presented in this study. As illustrated, similar fold change patterns were observed with a Pearson’s correlation coefficient of 0.74. Further analysis corroborated the effectiveness of our analysis on removal of batch effects (Fig S14). d) To ensure that the observed fold changes are not due to presence of some outliers in our samples, we performed jack-knife resampling. Repeating 100 times, we sampled from 20% to 90% of the quantile normalized transcriptome data from n=226 samples, while preserving the proportion of ASD-to-TD samples. The sampled datasets were then processed independently and fold change patterns were observed. As shown, we found that the jack-knife fold change patterns correlate well with those of total 226 samples as measured by Pearson’s correlation coefficient. This result demonstrates that the fold change patterns are shared in a large fraction of samples. We reached to similar conclusions based on the resampling of the network activity as presented in Fig. 6 and Fig. S11. e) To estimate the signal-to-noise ratio of observed fold changes on n=226 samples, we repeated the jack-knife resampling procedure, but this time counted the number of DE genes that were also identified as differentially expressed in the sampled datasets (limma analysis; FDR <0.05). As illustrated, we found that a large sample size is required to identify our DE list as significant. f-i) To assess the reproducibility of fold change patterns of differentially expressed genes, we performed another transcriptome experiment using a different microarray chip platform. The primary dataset was analyzed on Illumina BeadChip HumanHT-12, while this dataset was analyzed on Illumina BeadChip HumanWG-6. This second dataset was composed of 56 male toddlers (n= 35 ASD and 21 TD) that were shared with the primary dataset (technical replicate dataset). In a third dataset, we included an additional n= 48 samples including 24 independent ASD male toddlers, while 21 out of 24 TD samples overlapped with the primary dataset. These two groups of samples were processed separately to assess the reproducibility of results at both technical and biological levels. The latter would potentially hint on the penetrance level of the dysregulation signal in ASD population. As illustrated, we observed good reproducibility at technical (panels f and g) and biological (panels h and i) levels with Pearson’s correlation coefficients of 0.83 and 0.73, respectively. To assess if the overlapping TD samples in the partially overlapping dataset are driving the signal, we excluded the 21 overlapping TD subjects from the primary dataset (panels h and i). We further performed the transcriptomics analysis of an entirely independent cohort of ASD and TD male toddlers using RNA-Seq platform and reached to a similar conclusion (Fig S4). Boxplots represent the median (horizontal line), lower and upper quartile values (box), and the range of values (whisker), and outliers (dots).

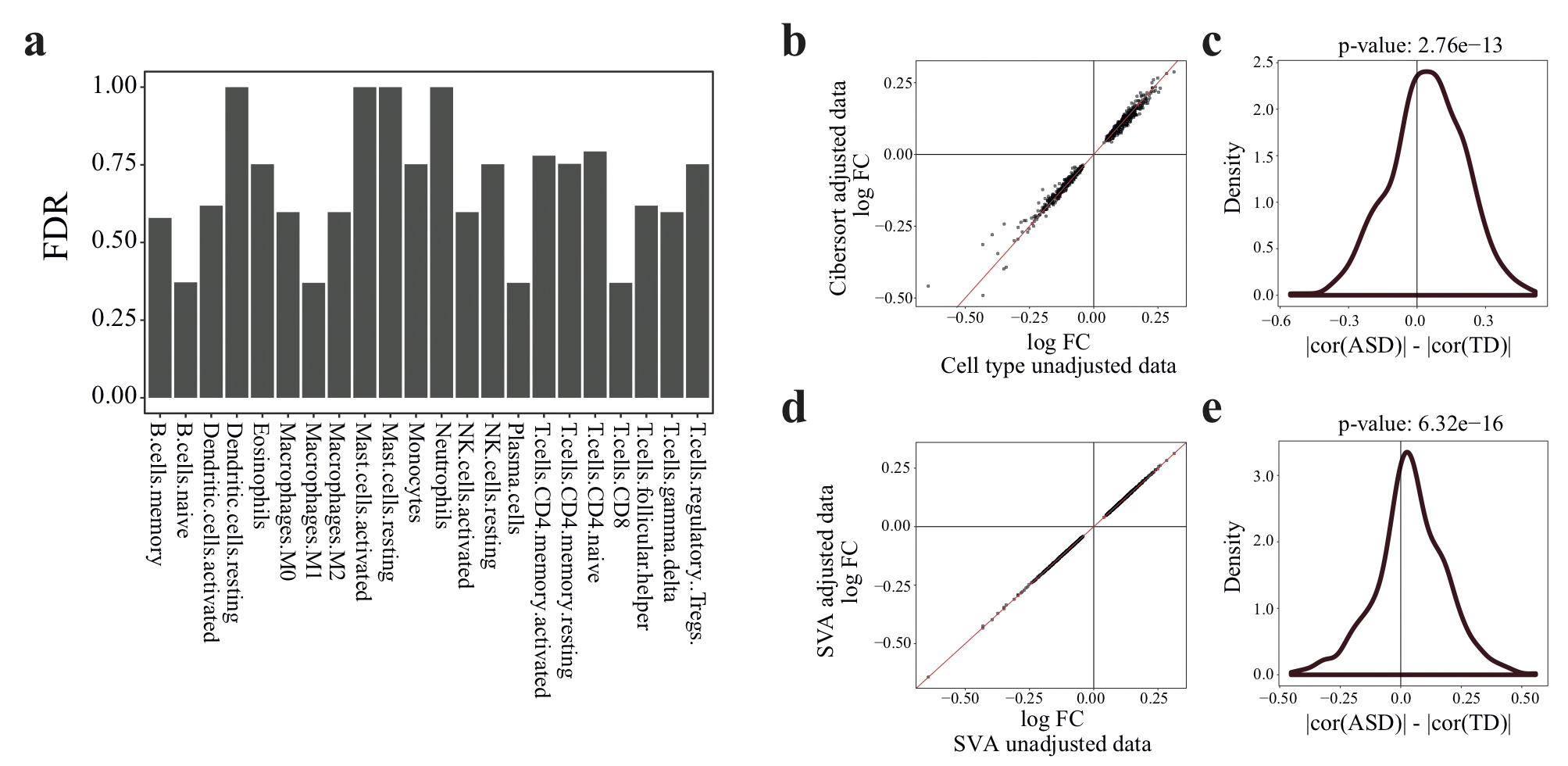

**Supplementary Figure 2**

Examining the presence of confounding factors in the gene expression data

**a)** Cell type compositions have not significantly changed between n=119 ASD and 107 TD toddlers. The cell type compositions were estimated in each sample using Cibersort algorithm. The relative frequency of each cell type was next compared between ASD and TD samples using t-test. p-values were adjusted using Benjamini-Hochberg procedure. **b)** To assess the potential confounding effect of cell type composition on the gene expression patterns, we included the cell types with nominal p-value <0.1 (four cell types) in the regression model. As illustrated, the fold change patterns remain robust to changes in the cell type composition. **c)** To assess the potential effect of cell type composition on the network activity, we regressed out the effect of the cell types that nominally changed between n=119 ASD and 107 TD (p-value <0.1) from the gene expression data. As illustrated, the DE network remains transcriptionally over-active. For an unbiased analysis, as is done in all comparisons of network activity between leukocyte ASD and TD samples, a merged network composed of union of interactions between DE-ASD and DE-TD networks were considered. The signal becomes stronger if the analysis was based on only DE-ASD network. Paired one-sided Wilcoxon-Mann-Whitney test used for the comparison of Pearson’s correlation coefficients. **d)** To examine potential effect of hidden confounding effects on the gene expression patterns, we performed SVA analysis. SVA predicted only one significant surrogate variable (SV) in the normalized but unadjusted microarray dataset (num.sv() function). The predicted SV was highly correlated with the experimental batch. We have already included the batch as a covariate in our analyses and have regressed it out prior to the calculations of correlation and mutual information scores. Accordingly, SVA did not find any significant SV in the batch adjusted expression data. To further explore the potential influence of other hidden confounding variables, To further explore the potential effect of (hidden) confounding co-variates on our results, we performed SVA on the batch adjusted dataset and included the first SV in the regression models related to the DE analysis (total of two covariates, one batch and one SV). As illustrated in Fig. S2c, we observed that the fold change patterns remained consistent. **e)** To assess the effect of hidden surrogate variables on the network activity, we considered the first SV from above as an additional covariate and regressed out its effect from gene expression data from n=119 ASD and 107 TD samples. As illustrated, the DE network remains transcriptionally over active (paired one-sided Wilcoxon-Mann-Whitney test).

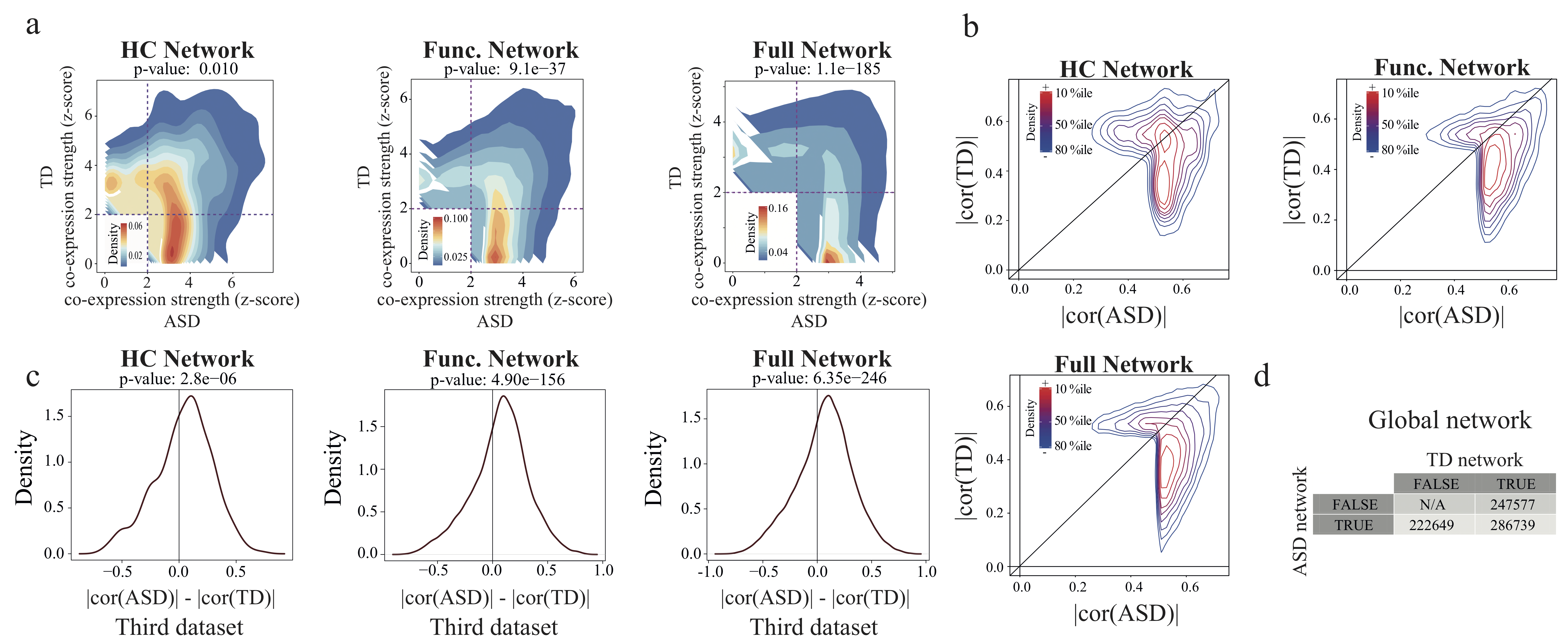

**Supplementary Figure 3**

Robustness analysis related to co-expression activity of DE network in ASD samples

a) The over-activity of the DE-ASD network is independent of the backbone static network. Here, the co-expression strength of context specific DE-ASD and DE-TD networks were compared using three different backbone networks of high confidence (HC), functional, and full co-expression across n=119 ASD and 107 TD samples (see methods for details). The three networks varied in number of genes and interactions. For each backbone, only those interactions that were significantly co-expressed (FDR <0.05) in at least one of diagnosis groups were included in the analysis. Red and blue colors represent regions with high and low density of interactions, respectively. The interaction strengths were compared by paired two-sided Wilcoxon-Mann-Whitney test. b) The over-activity of the DE-ASD network was uncovered using a mutual information-based method. To assess whether the elevated co-expression strength of the networks are supported with other metrics, we calculated the Pearson’s correlation coefficient for each interaction present in the static networks or all possible pairs of DE genes in the case of full co-expression network (n=119 ASD and 107 TD subjects). We next only retained interactions that had an absolute correlation of above 0.5 in either ASD or TD diagnosis groups. As illustrated, interactions tend to have higher correlations in ASD than TD, indicating the robustness of observed over-activity across different similarity metrics. c) The over-activity of the DE network was examined in another dataset that included n=24 independent male ASD toddlers. This dataset also contained n=24 TD male toddlers, including 21 subjects that overlapped with the primary dataset. As shown, we observed replicable over-activity of the DE networks in ASD group, as measured based on the co-expression strength. The correlation strengths of interactions in ASD and TD samples were compared using paired two-sided Wilcoxon-Mann-Whitney test. We further performed the transcriptomics analysis of an entirely independent cohort of ASD and TD male toddlers using RNA-Seq platform and reached to a similar conclusion (Fig S4). d) To assess whether network over-activity of ASD samples is a general characteristic in our dataset or is specific to the DE networks, we constructed a global ASD network using the same approach as employed for the DE networks, but not limiting the network to the DE genes (n=119 ASD and 107 TD subjects). Thus, the backbone static network included all functional interactions present in GeneMania database. As shown, we found that in contrast to the DE networks, the global network showed slightly but significantly higher co-expression levels in TD samples (p-value 1.0; paired one-sided Wilcoxon-Mann-Whitney test). The table indicates the number of interactions in the global network that were deemed as significant in either ASD or TD diagnosis groups (FDR <0.05).

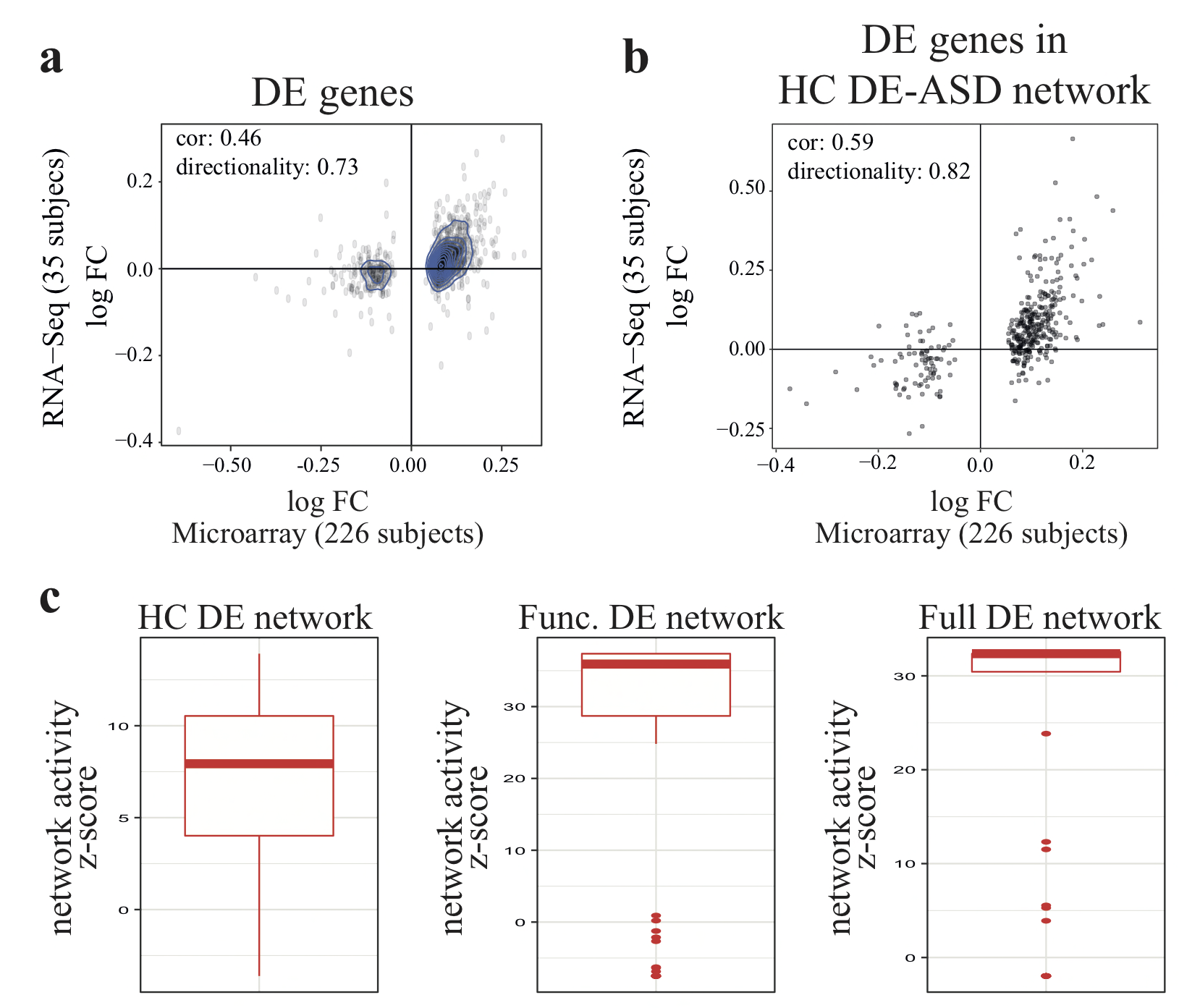

**Supplementary Figure 4**

Reproducibility of the signature in an independent cohort as measured by RNA-Seq

To assess the generalizability and reproducibility of the observed signature, we performed transcriptomics analysis of an independent cohort of male toddlers with n=12 TD and 23 ASD (56 samples including technical replicates, Table S1). Our primary dataset was analyzed by microarray platform. To ensure the results are not dependent on the transcriptomics platform, we analyzed this dataset using RNA-Seq platform. Moreover, we included in the dataset 7 and 14 technical replicated of TD and ASD samples. Technical replicates were modeled as random effects in the differential expression analysis of this RNA-Seq dataset. None of subjects overlapped with those in the primary dataset. **a)** Fold change comparison of the 1236 DE genes between the microarray and RNA-Seq datasets demonstrated moderate conservation of the fold change patterns between the two datasets (Pearson’s correlation coefficient: 0.46). We also observed 73% of DE genes preserved their directionality in both datasets (e.g., up or down in both datasets). The moderate conservation of the fold change patterns could be due to inherent heterogeneity within ASD samples (the two datasets included non-overlapping subjects), relatively small size of the RNA-Seq dataset, false positives among 1236 DE genes, or technical differences between the two transcriptomics platforms. **b)** Genes involved in the DE-ASD network exhibit highly preserved fold change patterns between the two datasets. The figure compares the fold change pattern of genes involved in HC DE-ASD network. We observed a boost in the Pearson’s correlation coefficient of the fold change patterns between the RNA-Seq (n=56 samples) and microarray (n=226 samples) datasets, suggesting the network construction procedure has removed some of the false positives among the 1236 DE genes. We also observed 82% of genes involved in the HC DE-ASD network have preserved their directionality between the two datasets. **c)** DE networks are transcriptionally over-active in the independent RNA-Seq dataset. Y-axis demonstrates the z-transformed p-value comparing the activity of DE network between ASD and TD samples in the RNA-Seq dataset. To ensure robustness of the results, iterating 100 times, we randomly selected n=12 samples from each diagnosis group (unique subjects) and compared the activity of DE network using a two-sided Wilcoxon-Mann-Whitney test. Y-axis demonstrates the z-transformed p-values. In cases that z-score could not be estimated (e.g., p-value =0), we used the z-score form lowest non-zero p-value. Boxplots represent the median (horizontal line), lower and upper quartile values (box), and the range of values (whisker), and outliers (dots).

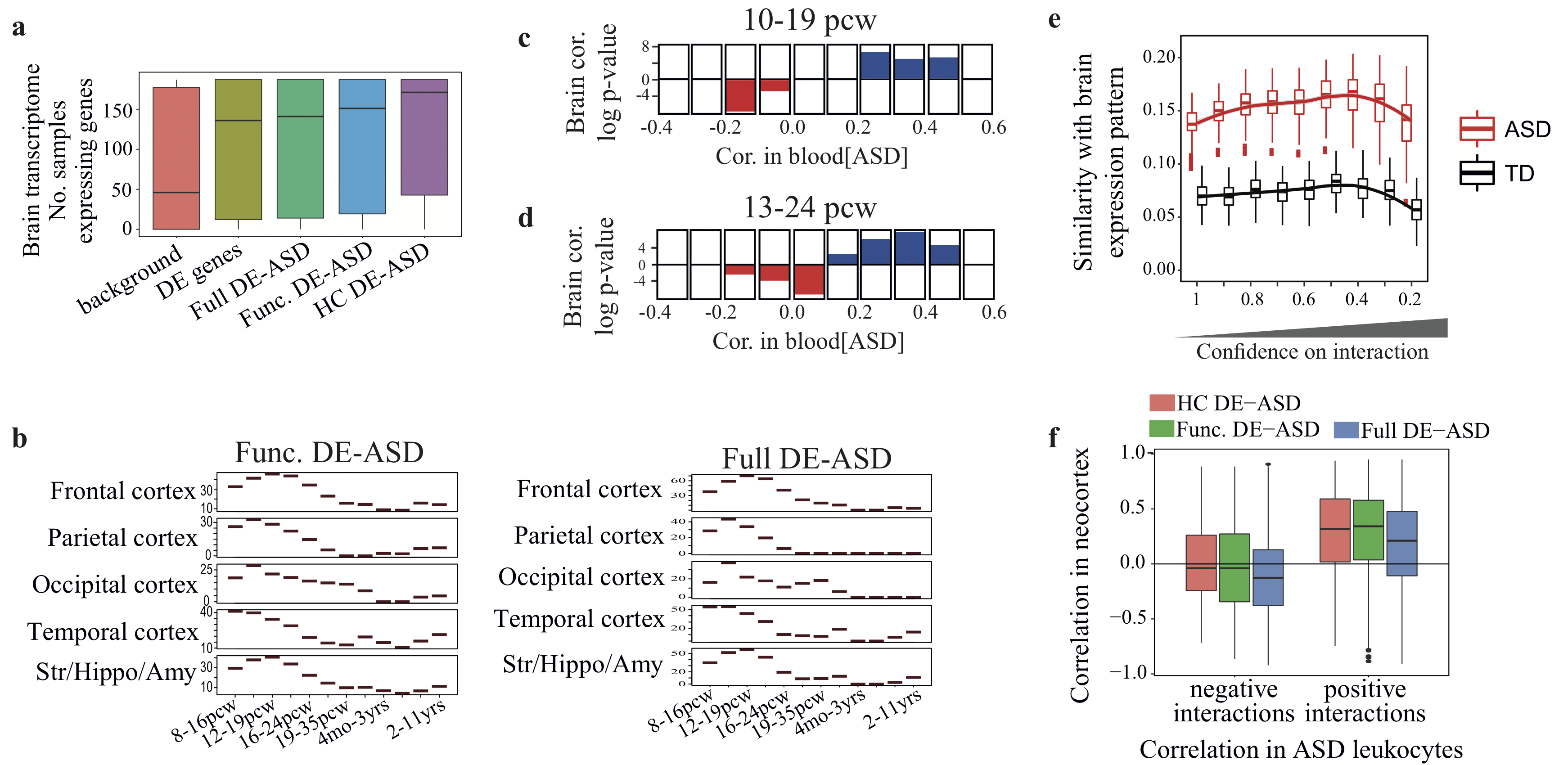

**Supplementary Figure 5**

DE genes are involved in networks that are preserved between blood and brain tissues

a) Genes involved in the DE-ASD networks are highly expressed during normal brain development process in prenatal (≥ 8 post conception weeks) and early postnatal (<1 year old) ages (p-value <5.2 x 10^-29^). BrainSpan RNA-Seq transcriptome data were used for this analysis (n=187 samples). Genes with RPKM >5 were considered as expressed in each sample. Groups were compared using two-sided Wilcoxon-Mann-Whitney test. b) Network activity patterns of functional and full DE-ASD networks based on BrainSpan transcriptome data at prenatal and early postnatal ages. At each time window, the activity was measured based on the distribution of co-expression strength of interacting gene pairs in DE-ASD network using Pearson’s correlation coefficient metric (n= 121 frontal, 73 temporal, 42 parietal, 27 occipital cortices, and 72 striatum, hippocampus, and amygdala samples across time points). The y-axis indicates the z-transformed p-value of co-expression strength as measured by a two-sided Wilcoxon-Mann-Whitney test. c) The conservation of interactions between blood and brain for a previously reported co-expression network around high confidence rASD genes in brain at 10–19 post conception weeks from Willsey et al. The interactions were partitioned based on their correlation value in the n=119 blood samples from subjects with ASD (window size of 0.1). The bar-graphs in each bin represents significant enrichment for positive (blue with positive enrichment values) or negative (red with negative enrichment values) interactions based on the corresponding brain transcriptome data (log_10_ transformed p-values; hypergeometric test). Only statistically significant (p <0.05) comparisons are represented in bar graphs. d) Similar to panel c, but for a co-expression network of high confidence ASD risk genes from 13 to 24 weeks post conception were extracted from Willsey et al. e) brain derived co-expression network of rASD genes (the same network as panel d) were compared to the co-expression pattern of the same interactions in leukocyte transcriptome of ASD and TD toddlers. Boxplots represent the observed similarity based on 100 random sub-sampling of n=75 ASD and 75 of TD samples (~70% of samples in each diagnosis group). The x-axis represents the top percentile of positive and negative interactions based on the brain transcriptome interaction weights. For example, 20 %ile illustrates the results when only top 20% of positively and top 20% of negatively interacting genes in the brain co-expression network were considered for the analysis (selected based on the interaction weights). As illustrated, ASD samples have significantly higher similarity in co-expression patterns with the developing brain than TD samples. f) Positive and negative interactions in DE-ASD networks were preserved between blood (n=119) and brain at prenatal and early postnatal ages (n=187). Boxplots represent the median (horizontal line), lower and upper quartile values (box), and the range of values (whisker), and outliers (dots).

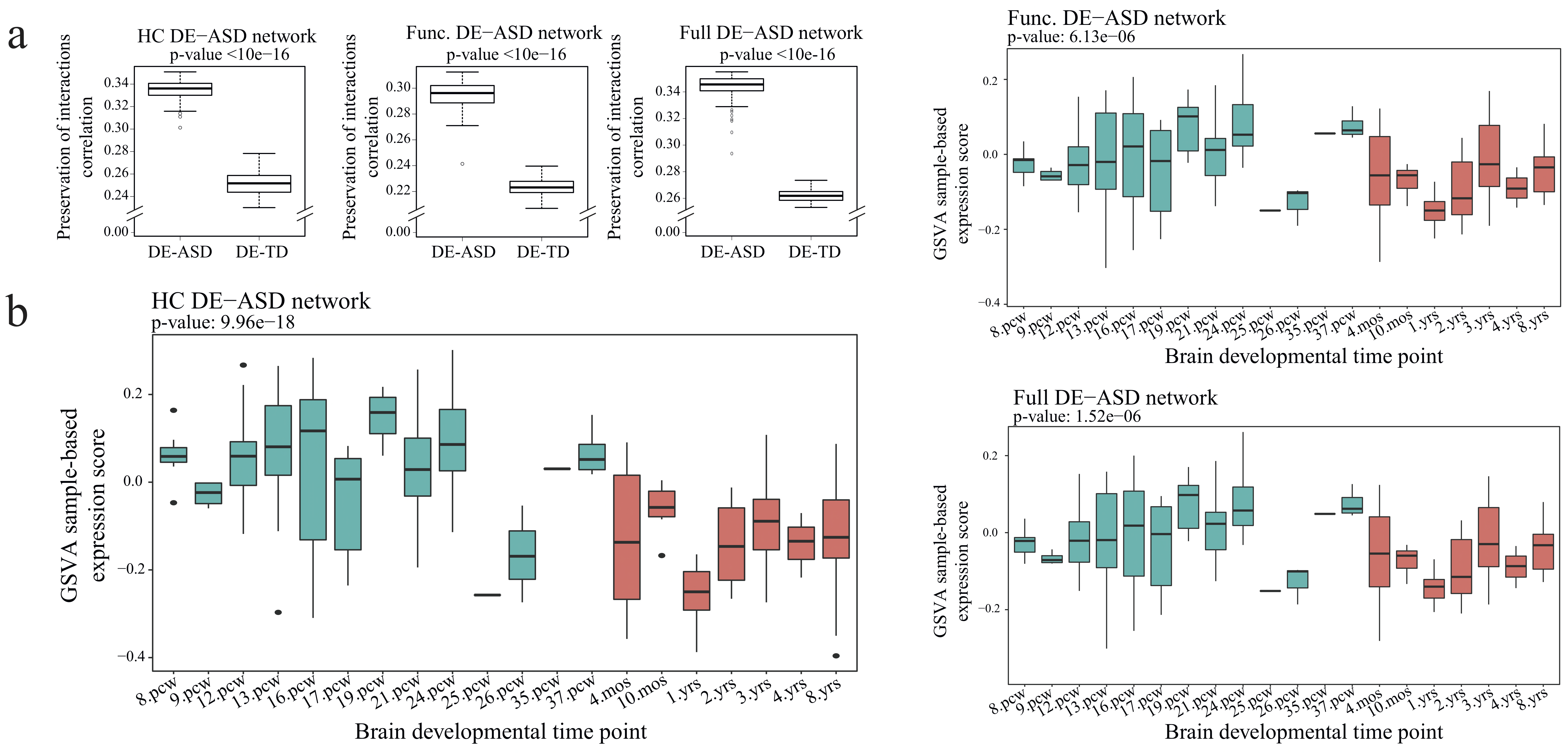

**Supplementary Figure 6**

DE-ASD network is transcriptionally active at prenatal brain

a) DE-ASD network is significantly better preserved than DE-TD network in prenatal and early postnatal brain. Correlations of interactions of DE-ASD and DE-TD with brain gene expression were estimated using BrainSpan RNA-Seq data. Briefly, to examine the preservation of interactions in each of the two networks, iterating 100 times, we calculated the correlations of interactions in DE-ASD and DE-TD networks based on a randomly selected subset of n=70 ASD and 70 TD samples, respectively. Next, the similarity of estimated correlations of interactions between brain and blood samples were calculated using Pearson’s correlation coefficient. As illustrated, the DE-ASD network is significantly better preserved in prenatal and early postnatal brain transcriptome data. b) Transcriptional over-activity of DE-ASD networks at prenatal brain development period. The transcriptional activity of genes involved in DE-ASD network were estimated in each sample using GSVA analysis. Opposed to our network transcriptional activity measure that is based on the co-expression magnitude of interactions, GSVA employs a sample based metric based on the concept of GSEA in which the overall expression pattern of the genes in each sample is examined, disregarding the network structure. As illustrated, similar to the co-expression-based analysis of network activity, GSVA supports up-regulation of DE-ASD networks at prenatal brain transcriptome, suggesting the robustness of the results to methodological variations. Reported p-values are based on the comparison of the DE-ASD network expression pattern between prenatal (n=157 samples) and early postnatal (n=90 samples; 4 month-old to 8 year-old) periods using a two-sided Wilcoxon-Mann-Whitney test. Boxplots represent the median (horizontal line), lower and upper quartile values (box), and the range of values (whisker), and outliers (dots).

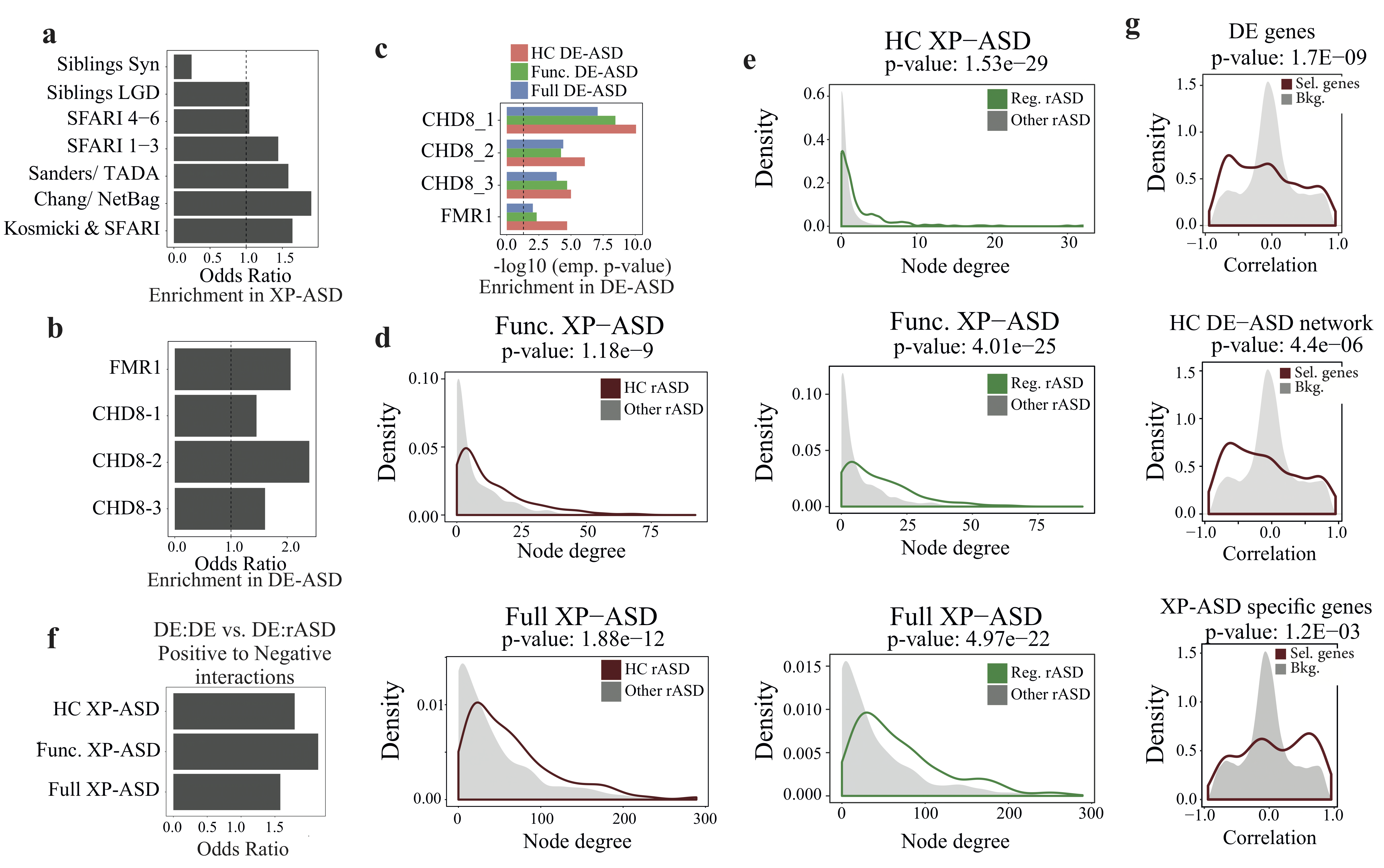

**Supplementary Figure 7**

Robustness analysis of observed association of rASD genes with DE-ASD networks

**a)** High confidence rASD genes are enriched in the XP-ASD networks. Genes with likely gene damaging (LGD) and synonymous (Syn) mutations in siblings of ASD subjects were extracted from Iossifov et al. study. **b-c)** The regulatory targets of well-known rASD genes (CHD8 and FMR1) are enriched in HC DE-ASD (panel b) as well as Functional and Full co-expression DE-ASD networks (panel c). The regulatory targets of *CHD8* were extracted from Sugathan et al. (*CHD8*-1), Gompers et al. (*CHD8*-2), and Cotney et al. (*CHD8*-3). The regulatory targets of *FMR1* gene were retrieved from Darnell et al. P-values were calculated empirically by permutation tests. **d)** High confidence rASD genes are more strongly associated with the XP-ASD network than the lower confidence ones, as judged with the number of interactions. The node degree distribution of high confidence and lower confidence rASD genes were compared using a two-sided Wilcoxon-Mann-Whitney test. **e)** rASD genes with potentially gene expression regulatory roles are enriched in the XP-ASD networks. The node degree of DNA binding rASD genes with those from the rest of rASD genes were compared using a two-sided Wilcoxon-Mann-Whitney test. **f)** Cross interactions between DE and rASD genes are significantly enriched for interactions with negative Pearson’s correlation coefficient, related to Fig 4. We compared the ratio of positive to negative interactions between DE and rASD genes to those within DE genes. The x-axis shows the estimated odds ratio. All p-values <3.1 x 10^-4^, two-sided Fisher’s exact test. **g)** Each plot shows the distribution of Pearson’s correlation coefficients of gene expressions with the time points during the in vitro differentiation process of primary human neural progenitor cells, related to Fig 4 (n= 77 samples; 3 fetal brain donors). As shown, DE genes are down-regulated during the differentiation process (negative correlations), while rASD genes show an up-regulation pattern (positive correlations).

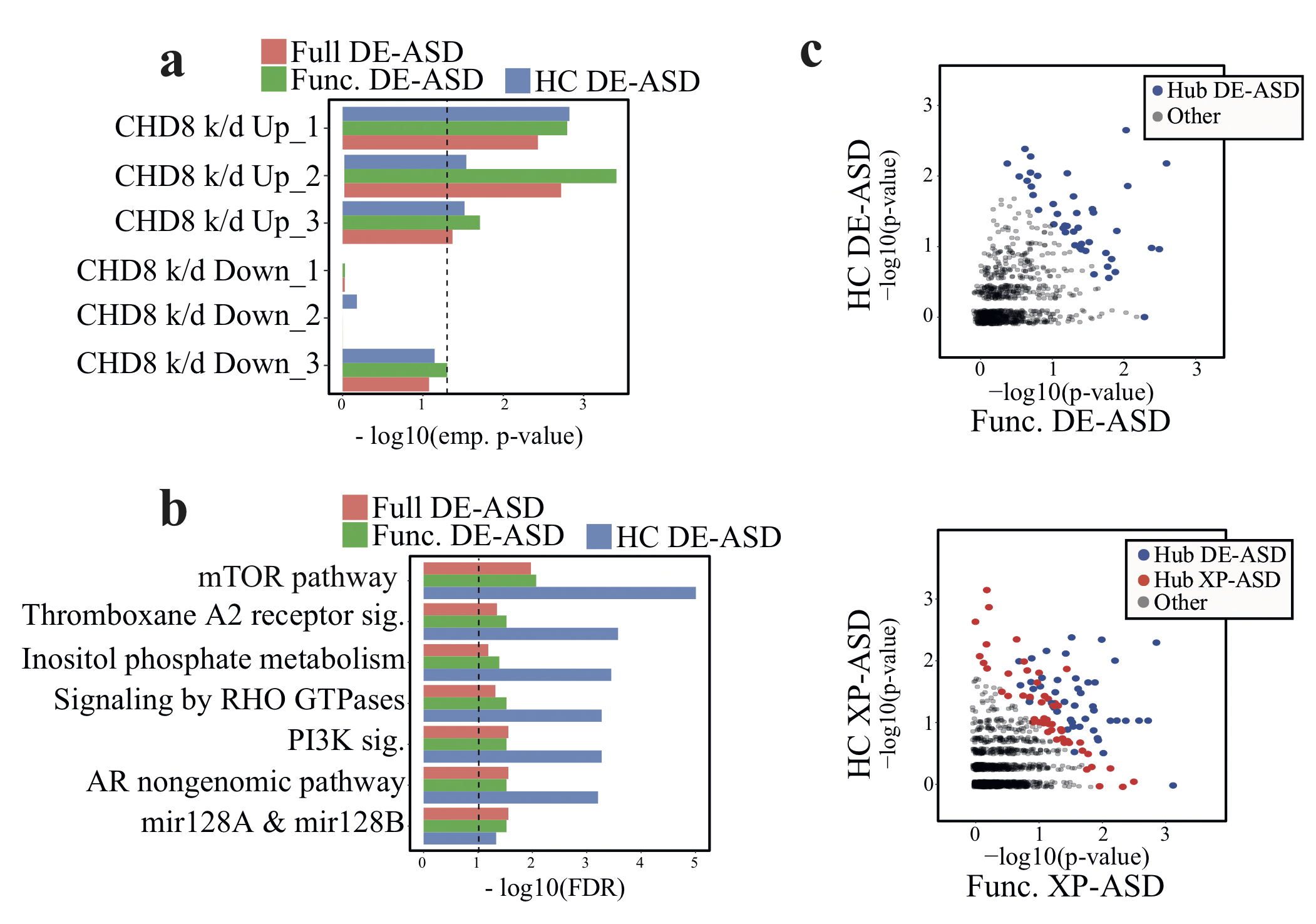

**Supplementary Figure 8**

Biological process enrichment analysis of the DE-ASD and XP-ASD networks

**a)** Genes up-regulated in background knock-down of *CHD8* are significantly enriched (permutation tests) in DE-ASD networks. Up and down regulated genes were extracted from Sugathan et al. (*CHD8*_1), Gompers et al. (*CHD8*_2), and Cotney et al. (*CHD8*_3). **b)** Biological processes that are enriched in the DE-ASD networks (Benjamini-Hochberg corrected FDR <0.1; hypergeometric test). The represented terms are also significantly changed between n=119 ASD and 107 TD samples as judged by GSEA. See methods for more details. **c)** Integrated hub analysis of DE-ASD and XP-ASD networks. For each network, the hub analysis was based on an integrated analysis of context specific high confidence (HC) and functional ASD networks. P-values are calculated empirically based on the degree distribution of genes involved in the DE-ASD and XP-ASD networks (see methods).

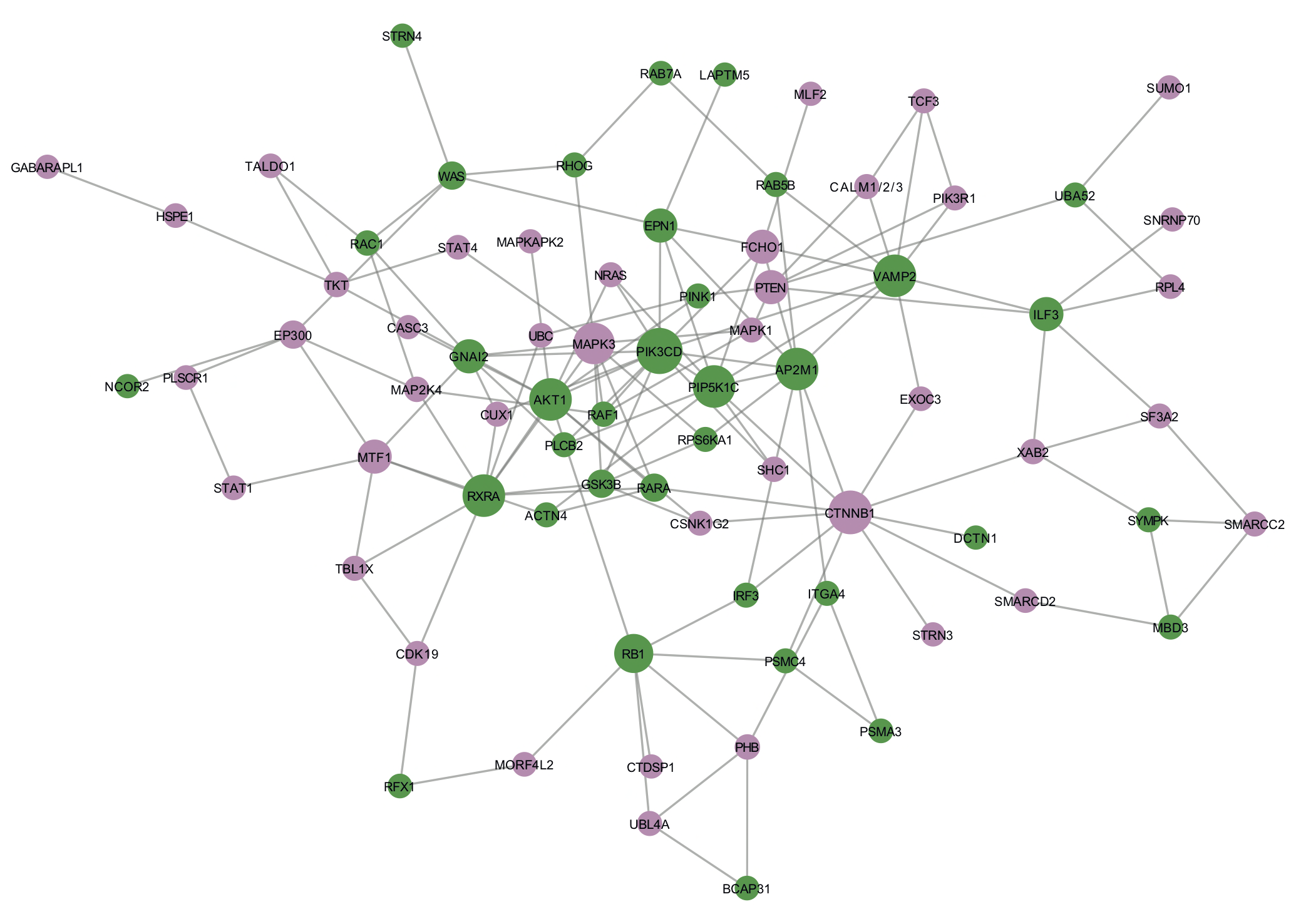

**Supplementary Figure 9**

Network of hub genes in the DE-ASD and XP-ASD networks

Network of hub genes in the HC XP-ASD network. Green color represents genes that are hub in both DE-ASD and XP-ASD networks. Purple color shows genes that are hub only in the HC XP-ASD network.

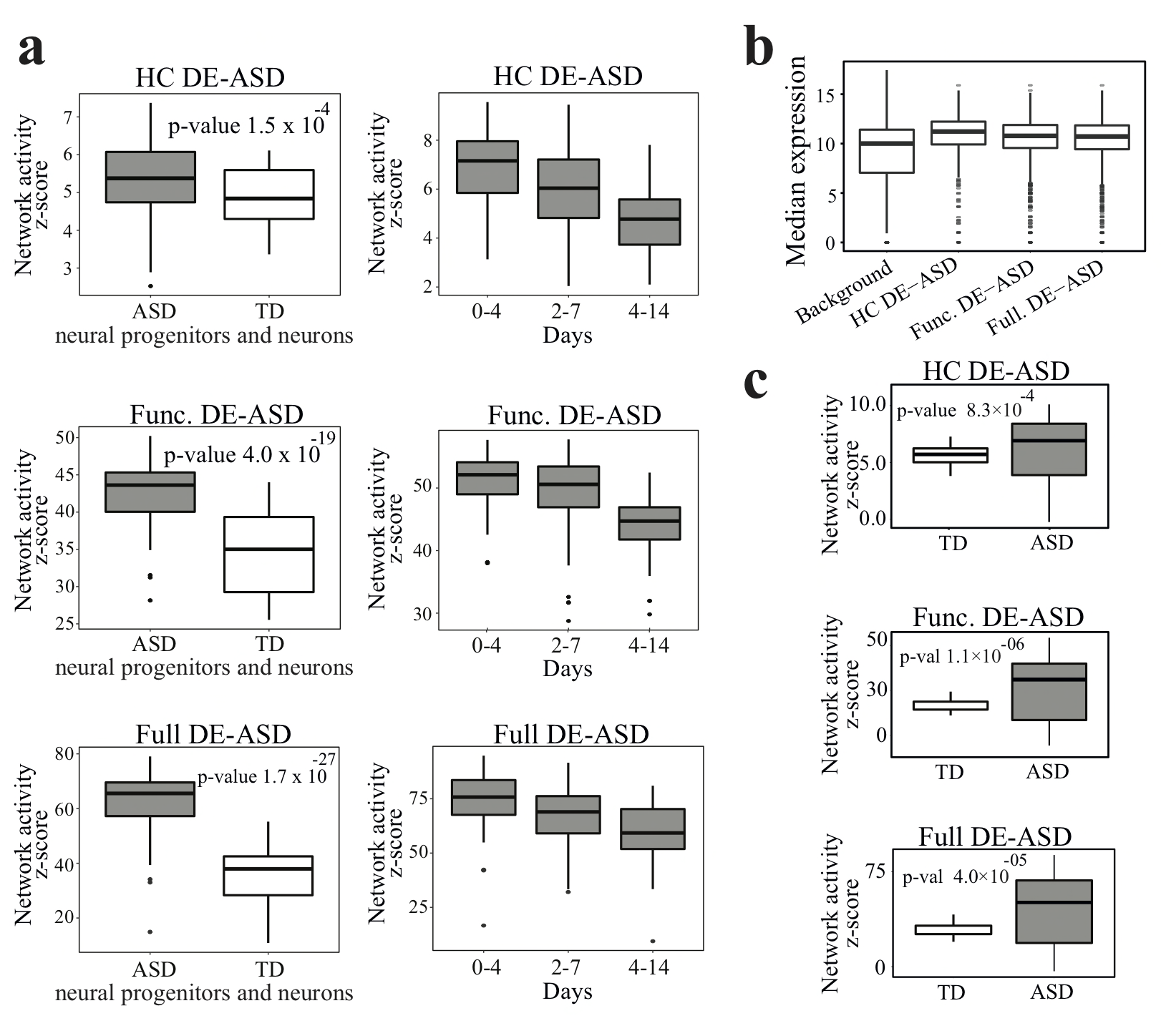

**Supplementary Figure 10**

Elevated co-expression of the DE-ASD networks in hiPSC-derived neuron models of subjects with ASD

a) DE-ASD networks are highly over-active in the ASD neural progenitor and neurons of individuals with ASD, compared to the TD cases. To ensure the robustness of estimated network co-expression activity levels, iterating 100 times, we measured the co-expression activity by randomly selecting 4 individuals within each diagnosis and measuring the co-expression strength across the neural progenitor to neuron stages of Day 0, 2, 4, 7, and 14 (n= 20 samples). The y-axis indicates the z-transformed p-value of co-expression strength as measured by two-sided Wilcoxon-Mann-Whitney test. b) The DE-ASD networks are highly expressed in RNA-Seq transcriptome data from hiPSC-to-neuron differentiation of individuals with ASD and TD from an additional dataset (GSE67528). In this dataset, the same set of hiPSC cell lines as those in the Fig. 6 are differentiated to neurons (in total three time points of hiPSC, NPC, and neurons) by an independent group (n= 83 samples from 8 individuals with ASD and 6 individuals with TD). RNA-Seq data were TMM normalized and log2(x+1) transformed. As shown, genes involved in the DE-ASD networks are highly expressed at NPC and neuron stages. c) The replication of observed dysregulation of DE-ASD network in the orthogonal dataset (GSE67528). To measure network activity, iterating 100 times, we randomly selected n=5 NPC and 5 neuron samples (in total 10 samples) from each diagnosis group. Boxplots represent the median (horizontal line), lower and upper quartile values (box), and the range of values (whisker), and outliers (dots).

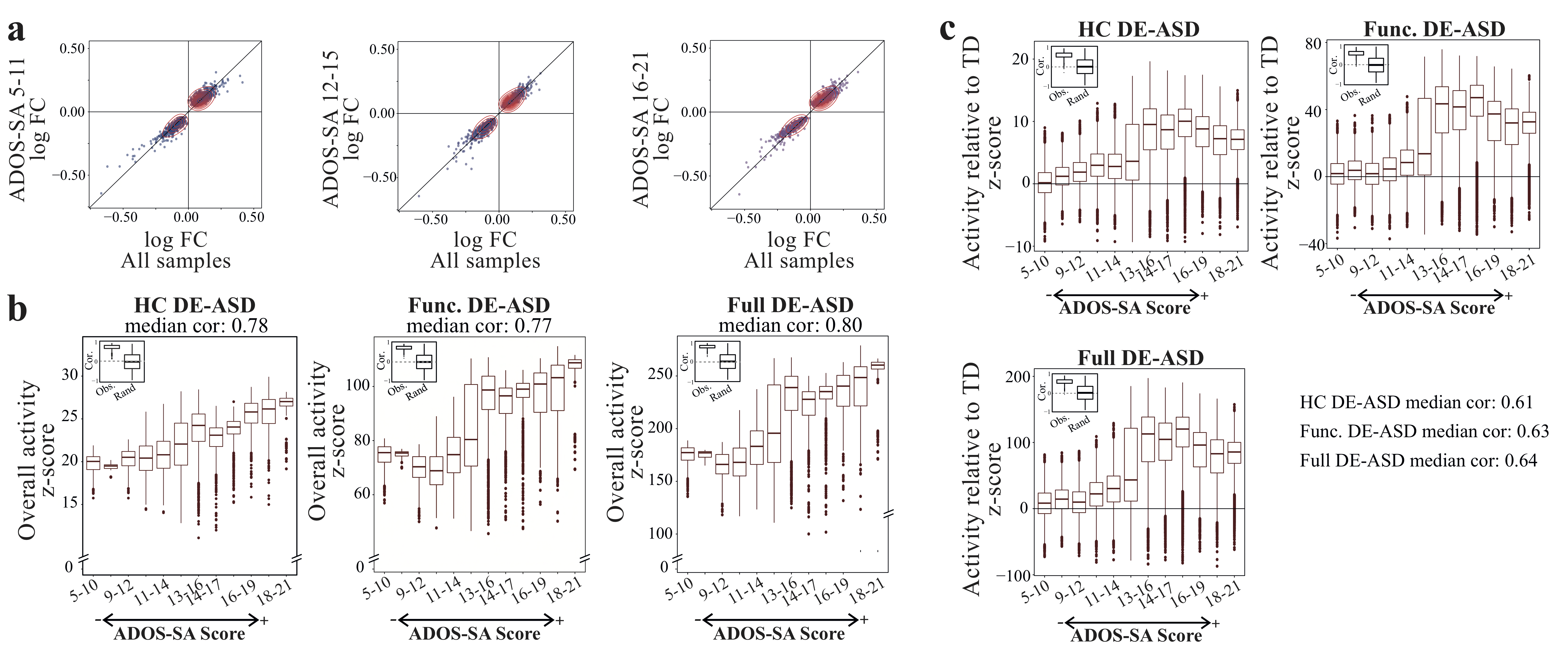

**Supplementary Figure 11**

The DE-ASD network transcriptional activity is correlated with ADOS-SA deficit scores

a) Male toddlers with ASD were categorized based on their ADOS social affect (ADOS-SA) deficit scores to the three groups of mild (ADOS-SA between 5 to 11; n=29), medium severity (ADOS-SA between 12 to 15; n=51), and high severity (ADOS-SA between 16 to 21; n=39). As shown, individuals at different severity levels show similar dysregulation patterns of DE genes. b) Male toddlers with ASD were sorted and grouped based on the ADOS-SA severity scores. Activity level of the DE-ASD networks in each group were measured based on the observed co-expression strength of interactions in the DE-ASD networks in randomly selected n=20 samples from each diagnosis group. The distribution of co-expression strengths (i.e., unsigned Pearson’s correlation coefficient) were next compared with what would be expected from a randomly selected set of genes within the same samples. The inset boxplots on top left demonstrate the distribution of observed and expected by chance Pearson’s correlation coefficient of ADOS-SA scores with network activity levels. The expected random distribution was generated by 10000 times random permutation of ADOS-SA scores of ASD individuals in the dataset. Note that the defined ASD severity levels are not independent and overlap with each other. c) The DE-ASD co-expression activity was measured by comparing the activity of DE-ASD networks in ASD versus randomly sampled TD cases (activity is measured based on n=20 selected samples from each diagnosis group). The inset boxplots demonstrate the distribution of observed and expected random Pearson’s correlation of ADOS-SA severity levels with the DE-ASD network activity levels. We used empirical methods to estimate the p-value of observed correlations with those of random in panels b and c. Iterating 10^6^ times, we sampled 100 data points from each of observed and random groups and assessed if the mean correlation in the samples from the random group is equal or higher than those of the observation group in absolute value. This analysis demonstrated a two-sided empirical p-value <10^-6^ for observed correlations. Boxplots represent the median (horizontal line), lower and upper quartile values (box), and the range of values (whisker), and outliers (dots).

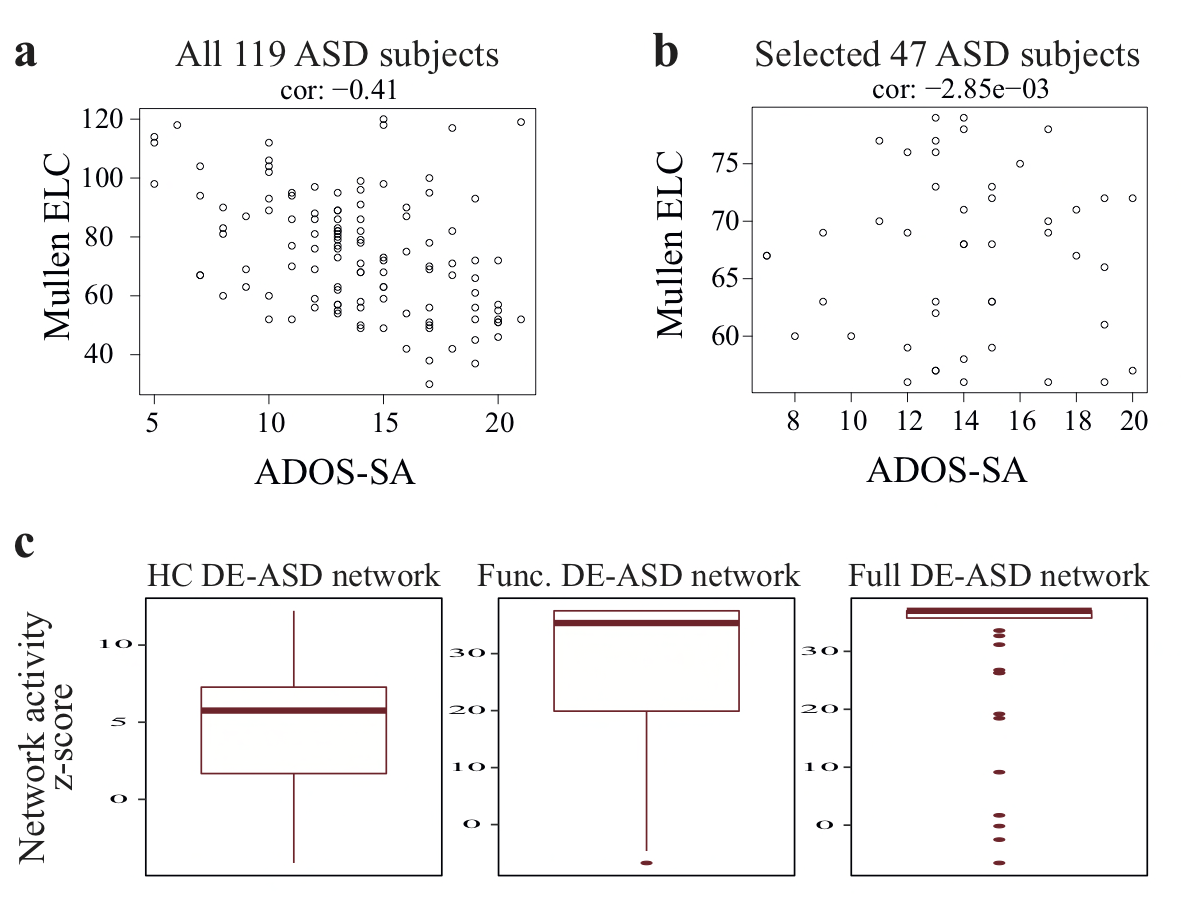

**Supplementary Figure 12**

Isolating the effect of ADOS-SA scores on the co-transcriptional activity of DE-ASD networks

**(a)** In our ASD cohort, ADOS social affect (ADOS-SA) scores are correlated with Mullen ELC scores (n=119 subjects with ASD; Pearson’s correlation coefficient: -0.41). **(b)** To isolate the effect of ADOS-SA scores, we selected subjects with ASD who have Mullen ELC scores above 55 and below 80 (n=47 subjects). In this subset, ADOS-SA scores were no longer correlated with Mullen ELC scores (Pearson’s correlation coefficient: -0.0029). We next divided the 47 ASD samples into two groups based on their median ADOS-SA scores. Iterating 100 times, we sampled n=15 from each ASD subgroup and compared the activity of the network. Panel **c** represents the z-transformed p-value of the comparisons of DE-ASD network over-activity between high and low ADOS-SA groups as measured by a one-sided Wilcoxon-Mann-Whitney test. In cases that z-score could not be estimated (e.g., p-value =0), we used the z-score form lowest none-zero p-value. As illustrated, the DE-ASD network exhibits over-activity in subjects with high ADOS-SA scores in this selected subset. Boxplots represent the median (horizontal line), lower and upper quartile values (box), and the range of values (whisker), and outliers (dots).

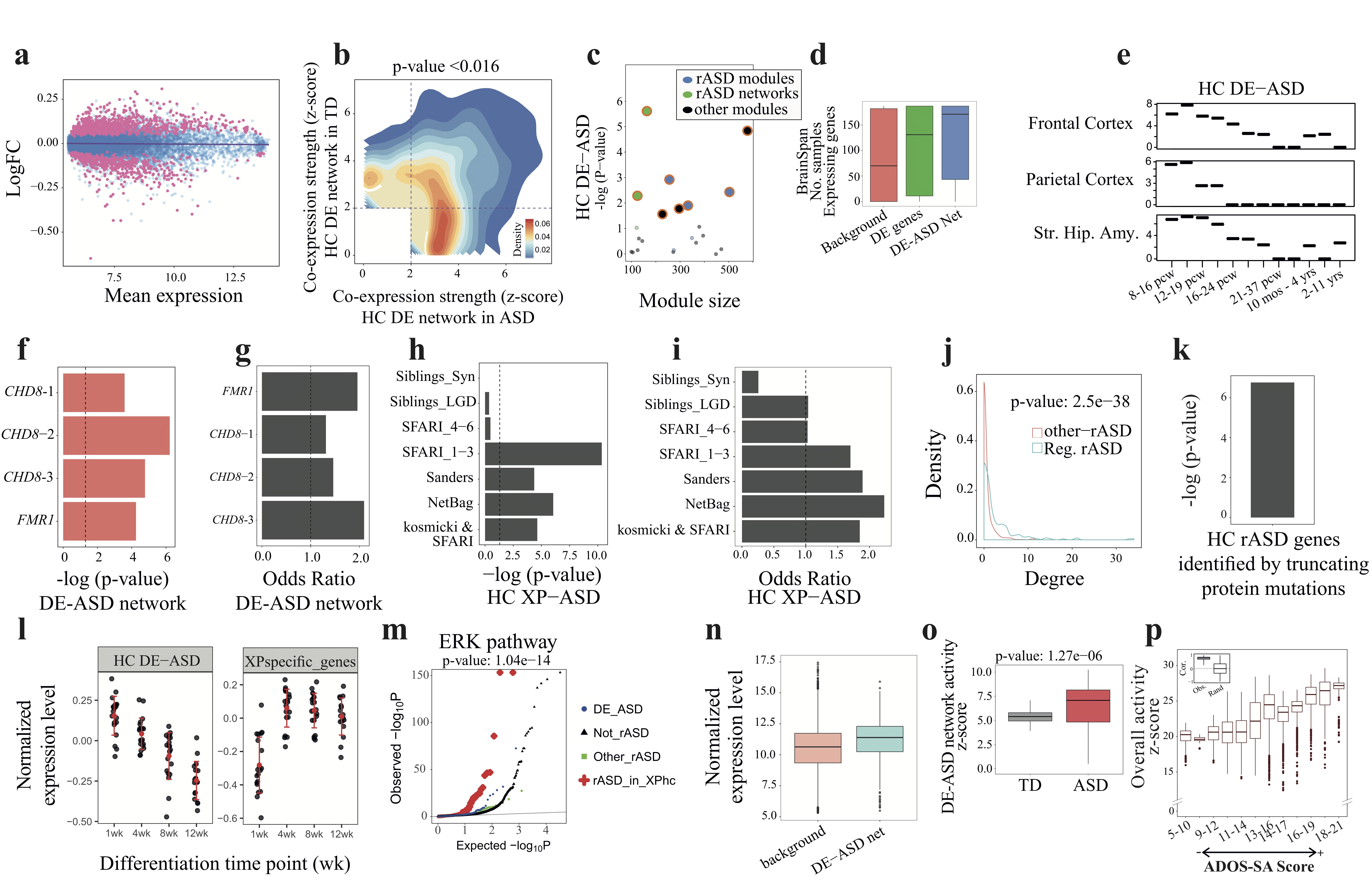

**Supplementary Figure 13**

Reproducibility of results under a different analysis setting

To assess the robustness of results, we assessed their reproducibility under a more stringent criterion for the expressed genes. The main results are based on the selection of 14,854 protein coding genes as expressed in the dataset. By comparison with the results of GTEx whole blood transcriptome data, we showed that number of expressed protein coding genes are in the same range in both studies (14,555 Expressed protein coding genes in GTEx; see methods for more details). Here, we employed a more stringent analysis by setting the p-value detection cut-off threshold at 0.01 instead of 0.05. This resulted in the selection of 13,032 protein coding genes as expressed in our leukocyte transcriptome dataset (n=119 ASD and 107 TD). We re-analyzed the new filtered dataset from DE analysis onward (constructed HC DE and XP networks). As shown, (a) MA-plot shows a similar pattern with mean expression being, in overall, uncorrelated with the fold change patterns. (b) the DE network showed transcriptional over-activity in ASD subjects (paired Wilcoxon-Mann-Whitney test). (c) The DE-ASD network significantly overlaps with the same modules and networks of rASD genes as the main results (permutation test). (d) The DE-ASD network is preferentially expressed at prenatal brain (n=187 BrainSpan neocortex samples). (e) Transcriptional activity of the DE-ASD network shows a peak at 10-19pcw in prenatal brain (z-transformed p-values of a two-sided Wilcoxon-Mann-Whitney test). (f-g) The DE-ASD network is significantly enriched for targets of high confidence rASD gens (permutation test). *CHD8*-1: Sugathan et al., *CHD8*-2: Cotney et al., *CHD8*-3: Gompers et al., *FMR1*: Darnell et al. (h-i) The XP-ASD network is preferentially associated with high confidence rASD genes (hypergeometric test). (j) The XP-ASD network is enriched for rASD genes with regulatory roles (two-sided Wilcoxon-Mann-Whitney test). (k) The high confidence rASD genes identified by truncating protein mutations in their sequence and pLI score >0.9 through large-scale genetics studies are enriched in the XP-ASD network (hypergeometric test). (l) The DE and rASD genes show anti-correlated expression patterns in in vitro neural differentiation data (n=77 samples from 3 fetal brain donors). (m) The XP-ASD network preferentially incorporates rASD genes with regulatory roles on RAS/ERK, PI3K/AKT, and WNT/B-catenin proteins. Results for RAS/ERK pathway is only shown due to space limitations; similar patterns were observed for PI3K/AKT and WNT/B-catenin pathways. (n) The DE-ASD network is preferentially expressed in hiPSC-derived ASD neurons and neural progenitor cells (two-sided Wilcoxon-Mann-Whitney test; n= 83 samples from 14 donors). (o) The DE-ASD network is significantly over-active at transcriptional level in ASD neural progenitor and neuron models (two-sided Wilcoxon-Mann-Whitney test; n= 5 progenitor and 5 neuron samples from each of ASD and TD groups). (p) Co-transcriptional activity of the DE-ASD network correlates with ADOS social affect scores of ASD subjects (permutation test; n=20 subjects at each ASD symptom severity level). Boxplots represent the median (horizontal line), lower and upper quartile values (box), and the range of values (whisker), and outliers (dots).

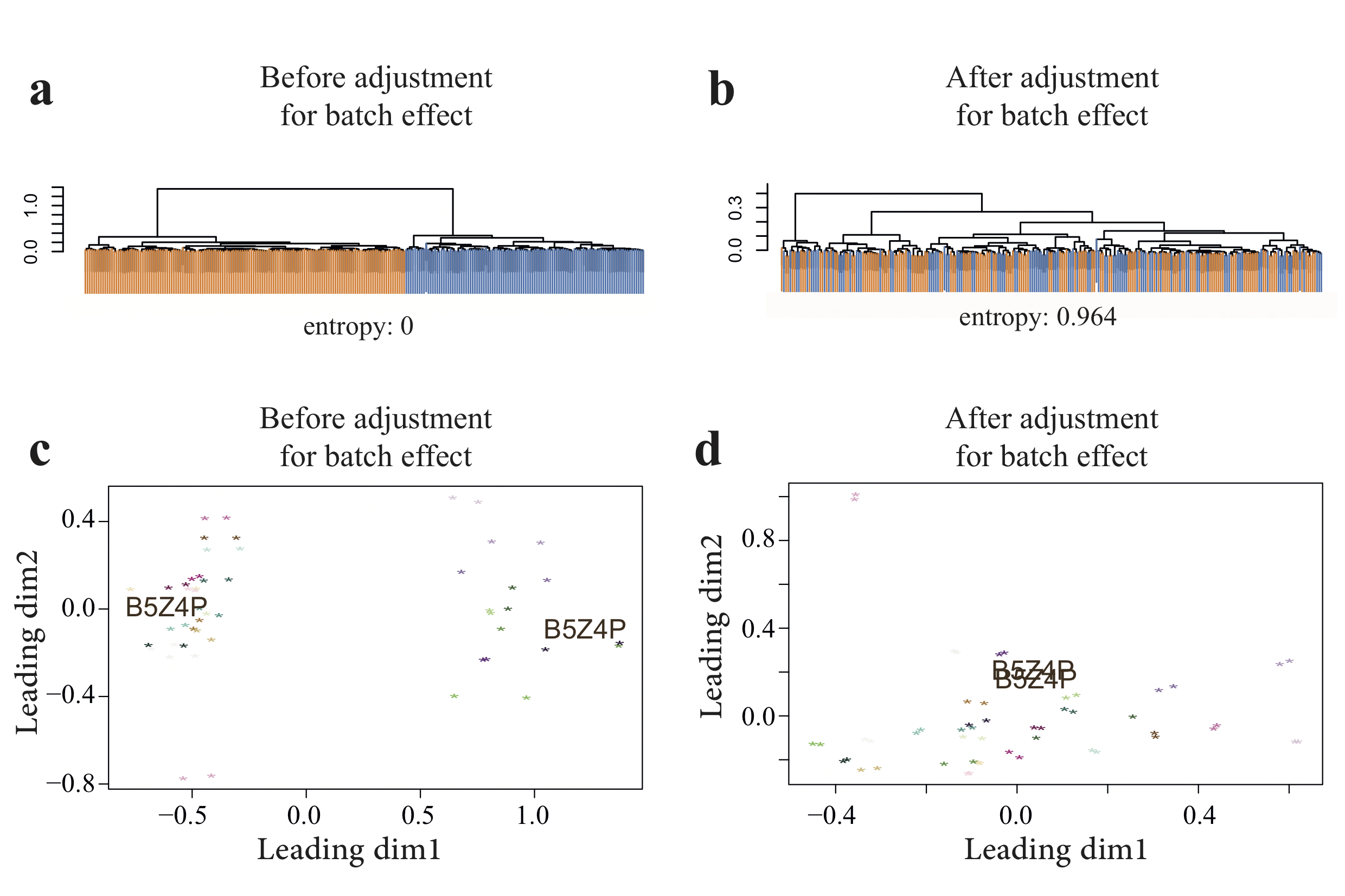

**Supplementary Figure 14**

Batch effects could be effectively handled by linear regression models

The top plots (a-b) illustrate a hierarchical clustering of n=226 subjects based on 1000 most variable genes in the primary microarray dataset. Samples are color coded based on the batch number. To estimate batch effects, we clustered the samples to 8 clusters and measured the uncertainty (heterogeneity) level of batch co-variate within each cluster using an entropy metric. The overall entropy score is sum of cluster entropy scores weighted by the cluster size. Entropy of zero implies that samples of the same batch are clustered with each other, and increasing entropy levels indicate a more random distribution of samples in terms of the batch co-variate. The bottom plots (c-d) demonstrate principle coordinate plot of samples that have technical replicates (57 samples). The distance of every two samples on the plot approximate their Euclidian distance based on 1000 most variable genes in the dataset. Samples are color-coded with technical replicates in the same color. B5Z4P is a sample with technical replicates in two different batches as obvious in panel (c). Principal component figures are generated by limma package in R.
